# Microbial byproducts determine reproductive fitness of free-living and parasitic nematodes

**DOI:** 10.1101/2021.08.02.454806

**Authors:** Mericien Venzon, Ritika Das, Daniel J. Luciano, Julia Burnett, Hyun Shin Park, Joseph Cooper Devlin, Eric T. Kool, Joel G. Belasco, E. Jane Albert Hubbard, Ken Cadwell

## Abstract

A distinguishing feature of *Trichuris* nematodes is that these parasitic worms reproduce within the digestive tracts of humans and other mammalian hosts shedding thousands of eggs daily, facilitating their sustained presence in the environment and hampering eradication efforts. Although this aspect of the lifecycle places *Trichuris* in a microbiota-rich environment, metabolic byproducts of bacteria that facilitate the reproductive development of parasites are unknown. Here, we employ a pipeline using the well-characterized free-living nematode *C. elegans* to identify microbial factors with conserved roles in the reproduction of nematodes. A screen for *E. coli* mutants that impair *C. elegans* fertility identified genes in fatty acid biosynthesis and ethanolamine utilization pathways, including *fabH* and *eutN*. *Trichuris muris* eggs displayed defective hatching in the presence of *E. coli* deficient in *fabH* or *eutN* due to reduction in arginine or elevated levels of aldehydes, respectively. Remarkably, *T. muris* reared in gnotobiotic mice colonized with these *E. coli* mutants displayed profound abnormalities including morphological defects and a failure to lay viable eggs. These findings indicate that microbial byproducts mediate evolutionarily conserved transkingdom interactions that impact the reproductive fitness of distantly-related nematodes.

## Introduction

The reproductive success of the parasitic nematode *Trichuris trichiura* is evidenced by the over 400 million individuals colonized by this soil-transmitted helminth (Vos et al., 2016). Its enormous reproductive capacity contributes to its infectious spread and hampers parasite eradication efforts. A distinct feature of *Trichuris* species is that reproductive development, from egg hatching to egg laying, is completed exclusively within the host digestive tract (Fahmy, 1954; Panesar, 1989). A new infection begins when embryonated eggs ingested from the environment hatch in the host cecum, a region within the gastrointestinal tract populated by a dense community of bacteria. Although bacteria mediate egg hatching for *Trichuris muris* (Hayes et al., 2010; Koyama, 2016; White et al., 2018), the *Trichuris* species that infects mice, a role for bacteria at later stages of *Trichuris* development remains unclear. Our group and others have shown that *Trichuris* infections in mice and humans affect the bacterial composition of the gut microbiota (Holm et al., 2015; Houlden et al., 2015; Jenkins et al., 2017; Lee et al., 2014; Ramanan et al., 2016; Rosa et al., 2021; Schachter et al., 2020; White et al., 2018). Additionally, the *T. muris* microbiota harbors taxa similar to those observed in the murine host microbiota, with a notable enrichment of Proteobacteria, including Gram-negative commensals like *E. coli* (White et al., 2018).

For the free-living nematode *C. elegans*, reducing the quantity or quality of the bacterial diet, or interfering with nutrient uptake, can impair germline development with consequences for fertility (Dalfo et al., 2012; Gracida and Eckmann, 2013; Korta et al., 2012; Pekar et al., 2017; Spanier et al., 2018; Starich et al., 2020). Because of the high degree of conservation of body plan and neuromuscular system organization within the phylum Nematoda, *C. elegans* has been used as a model for the discovery of anthelmintics for more than four decades (Salinas and Risi, 2018; Sepulveda-Crespo et al., 2020). Therefore, we hypothesized that bacteria-derived essential requirements for reproductive development in the well-characterized, model organism *C. elegans* are conserved in *Trichuris*.

## Results

### High-throughput screen identifies *E. coli* mutants that delay *C. elegans* fertility

To identify bacteria-derived factors that impact nematode reproduction, we assessed fertility of young adult *C. elegans* raised in 96-well format on a library of viable *E. coli* mutants suspended in liquid media. We used *C. elegans* bearing a temperature-sensitive mutation in the gene encoding the GLP-1 Notch receptor, *glp-1(e2141),* to facilitate quantification of fertility. At the semi-permissive temperature of 20°C, the number of germline progenitor cells is roughly half of the wild type (Michaelson et al., 2010), thereby sensitizing the worms to fertility defects when signaling pathways conveying nutritional sufficiency are altered (Dalfo et al., 2012; Korta et al., 2012). The *C. elegans* strain also carried fluorescent markers for the pharynx and for embryos (Roy et al., 2018) to facilitate counting and staging the worms, and to determine whether individual worms contained embryos (gravid) (Fig. 1, A-B). Wells were excluded from subsequent analyses if *E. coli* failed to grow, or if *C. elegans* exhibited developmental arrest or severely delayed growth. Using a series of selection criteria that considered both plate-by-plate statistical comparisons and penetrance, our primary screen of ∼3000 of the 3985 *E. coli* mutants in the Keio library identified 315 bacterial mutants that reduced the percentage of gravid worms at the established time point (Fig. 1C).

**Fig. 1.**
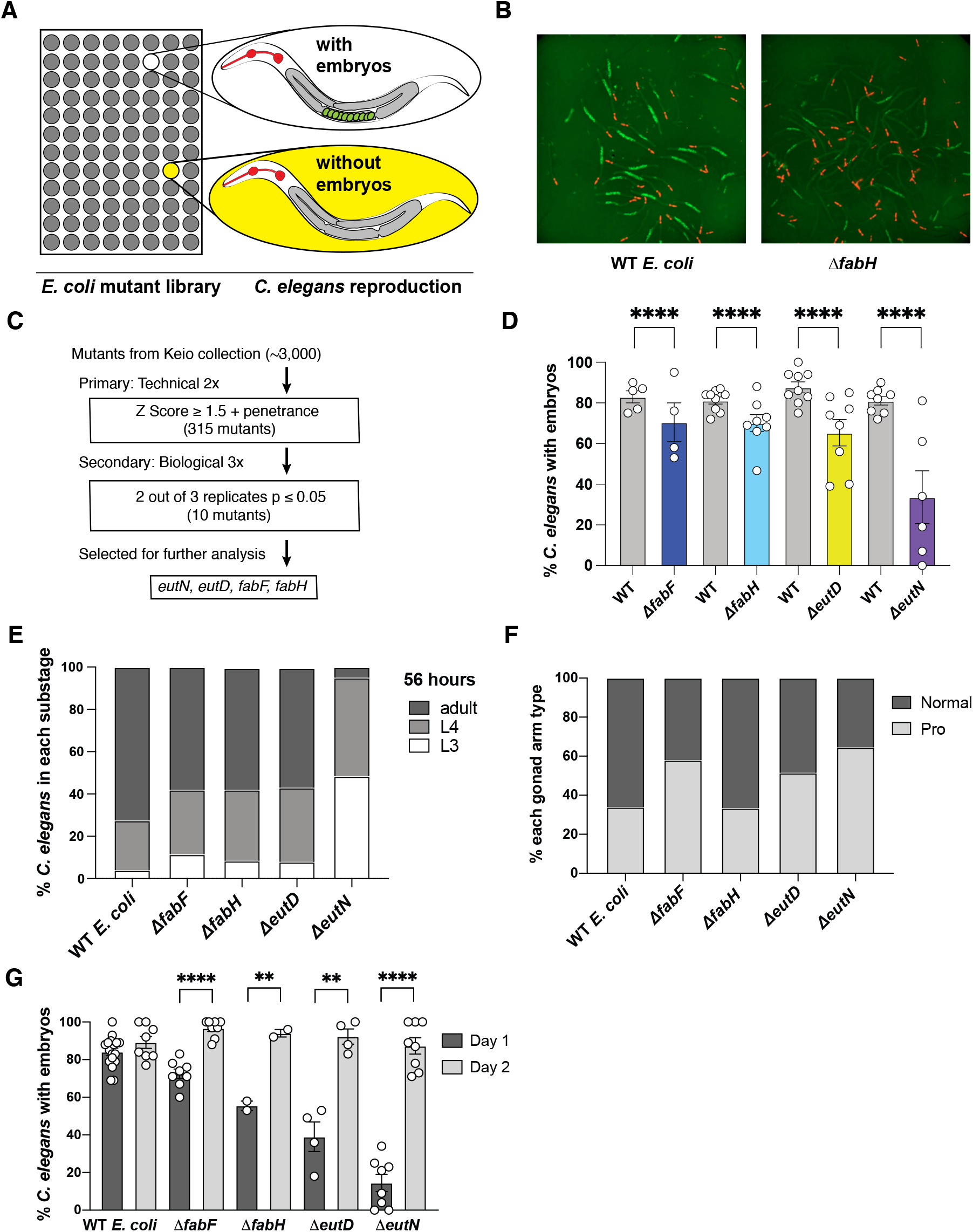
Identification of *E. coli* mutants that delay *C. elegans* fertility. **(A)** Concept of high throughput screening of *C. elegans* raised on *E. coli* mutants from the Keio library. **(B)** Example of reduced penetrance of *C. elegans* bearing embryos at a single time point; embryos marked with GFP and pharynx with mCherry. **(C)** Detailed flowchart for schematic in (A). **(D)** Percentage of *C. elegans* bearing embryos in replicates of four *E. coli* mutants of interest from the screen. Fisher’s exact test for each mutant versus controls from the same plates; total n number of worms scored are (left to right): 913, 411, 1253, 583, 1643, 550, 1561, 232. **(E)** Somatic development is delayed in *C. elegans* raised on *ΔeutN*. Percent of worms at indicated stage at 56 hours after L1 seeding to wells containing the control or mutant bacteria. n ≥ 38 worms were scored for each *E. coli* mutant. *ΔeutN* is the only mutant that is statistically significantly different from the control GC1547 (p ≤ 0.001). n number of worms scored (left to right): 52, 59, 61, 40, 54. **(F)** *ΔeutN* and *ΔfabF* delay germline development relative to somatic development. Percent of gonad arms displaying characteristic Pro phenotype. p ≤ 0.001 for *ΔeutN* and *ΔfabF*. Pooled from two independent trials; n number of worms scored are (left to right): 87, 103, 79, 41, 61. **(G)** Fertility is delayed. Percent of worms with embryos on Day 1 and Day 2. p ≥ 0.05 (not significant) for any mutant versus the control on Day 2. n number of worms scored (left to right): 547, 297, 242, 242, 60, 46, 124, 121, 253, 261. (D-G) Circles represent mean of independent biological replicates and bars show mean and SEM; pairwise Fisher’s exact test versus controls (D-F) or Day 1 versus Day 2 (G). ** p < 0.01, **** p < 0.0001.

After these 315 *E. coli* mutants were re-screened in biological triplicate, ten mutants significantly reduced the penetrance of embryo-bearing *C. elegans* compared to our wild-type (WT) *E. coli* control strain (GC1547) in at least two out of three biological replicates (Fig. 1C and S1, A-C). Seven of the ten *E. coli* genes fell into three functional groups related to ethanolamine utilization, fatty acid biosynthesis or lipopolysaccharide synthesis (Table S1). We focused on the *E. coli* mutants associated with the two functional groups fatty acid biosynthesis and ethanolamine utilization: *ΔfabF, ΔfabH, ΔeutD,* and *ΔeutN*. We pursued these functional groups because there were two mutants each that affected these processes and these mutants displayed consistent results across experiments, increasing our confidence that they represented bona fide hits (Fig. 1D). Complementation of the *ΔfabH*, *ΔeutN,* and *ΔeutD* mutations with a plasmid-borne copy of the corresponding gene confirmed their functional relevance in the *C. elegans* fertility delay phenotype (Fig. S1D-E). For *ΔfabF*, we rederived the *E. coli* deletion mutant by using P1 phage to transduce the original Δ*fabF*::*kan* allele from the Keio library strain into the Keio progenitor strain BW25113 (Fig. S1, F-G). Similar to the original mutant, the transductant caused a significant reduction in the percentage of embryo-bearing *C. elegans*.

To assess whether these *E. coli* mutants caused delays in somatic and/or germline development of *C. elegans*, we performed two different assays. First, we assayed somatic development using a time-course analysis of vulval development (Mok et al., 2015). Among the four mutants, only worms fed *ΔeutN* exhibited delayed somatic development (Fig. 1E). We also assayed for a delay in germline relative to somatic development (Fig. 1F) using as an assay enhancement of the temperature-sensitive Pro phenotype seen in a weak *glp-1(gf)* mutant (Pepper et al., 2003). Enhancement of Pro at the permissive temperature often occurs as a result of delayed germline development relative to somatic development (Hubbard and Schedl, 2019). Taken together, within the limits of these two assays we infer that the soma and germline of worms fed *ΔfabH* developed comparably with worms fed WT *E. coli, ΔfabF* and *ΔeutD* displayed delayed germline development without a somatic delay, and Δ*eutN* delayed somatic development as well as germline development relative to the (already delayed) somatic development.

Because our screen assessed the proportion of gravid *C. elegans* at a single time point, we asked whether non-gravid worms were permanently sterile or were delayed in achieving fertility. Exact time points corresponding to larval development differ when *C. elegans* is grown in liquid versus on solid media. Our screening time point (“Day 1” 62-65 hours post seeding; see Methods) is an early adult timepoint when worms fed on the WT bacteria are virtually all gravid (excluding a low background sterility of GC1474). When we prolonged the assay by scoring the worms on Day 2—24 hours after the initial screening time point—the percentage of gravid worms significantly increased, indicating that feeding on these *E. coli* mutants delayed *C. elegans* fertility rather than induced permanent sterility (Fig. 1G). We also found no significant differences when comparing the number of germline progenitor zone (PZ) nuclei in *C. elegans* raised on each mutant or WT *E. coli* at the L4-to-adult molt (Fig. S1, H-I), indicating that the delay in fertility is not likely due to slower accumulation of the PZ during larval stages (Agarwal et al., 2018; Kocsisova et al., 2019; Korta et al., 2012).

DAF-2 insulin/IGF-like and DAF-7 TGFß signaling pathways promote germline progenitor expansion and fertility in response to adequate nutrition, and this effect is also observed in the *glp-1* mutant background used in our screen (Dalfo et al., 2012; Michaelson et al., 2010; Pekar et al., 2017; Starich et al., 2020). Activation of these pathways ultimately interferes with the activity of downstream transcriptional regulators, *daf-16* and *daf-5*, respectively; loss of *daf-16* or *daf-5* therefore mimics constitutive activation of these pathways. We wished to determine whether loss of *daf-16* or *daf-5,* neither of which alter overall growth rate, would circumvent the fertility delay caused by each of the *E. coli* mutant diets. We fed WT or mutant *E. coli* to *daf-16* and *daf-5* mutant *C. elegans* and determined the percentage of gravid worms. Loss of *daf-16* actually exacerbated instead of rescuing the fertility defect of *C. elegans* raised on each mutant *E. coli*, as well as on WT *E. coli* (Fig. S1J). Although constitutive activation of the DAF-7 TGFß pathway via loss of *daf-5* partially and variably suppressed the fertility defect of *C. elegans* raised on *ΔeutN*, *ΔfabF* and *ΔfabH* (Fig. S1K), it also elevated the penetrance of gravid *C. elegans* raised on WT *E. coli* to a similar extent, making the suppression difficult to interpret. In short, we do not observe a simple causal relationship between either the insulin or TGFß signaling pathways as assayed by loss of *daf-16* or *daf-5*, and the alteration in the timing of *C. elegans* fertility caused by these four *E. coli* mutants.

### *E. coli* mutants with altered metabolic byproducts delay *C. elegans* fertility and inhibit *T. muris* egg hatching

Having identified *E. coli* mutants that delay fertility of a free-living nematode, we examined whether these bacteria impact a distantly related parasitic nematode. *T. muris* egg hatching was shown to be the reproductive stage sensitive to the presence of bacteria (Hayes et al., 2010; White et al., 2018). Thus, to begin investigating possible effects of the mutants identified in our *C. elegans* screen on the *Trichuris* lifecycle, we adapted a previously described method to quantify *E. coli*-mediated hatching kinetics of embryonated *T. muris* eggs *in vitro* (Hayes et al., 2010) (Fig. 2A). All *E. coli* mutants were confirmed to have similar growth kinetics to WT *E. coli* (Fig. S2A). Two of the mutants tested, *ΔeutN* and *ΔfabH,* elicited significantly reduced hatching compared to rates mediated by WT *E. coli* (Fig. 2B). Complementation of *ΔeutN* or *ΔfabH* with a plasmid-borne copy of the corresponding gene rescued hatching (Fig. S2, B-C).

**Fig. 2.**
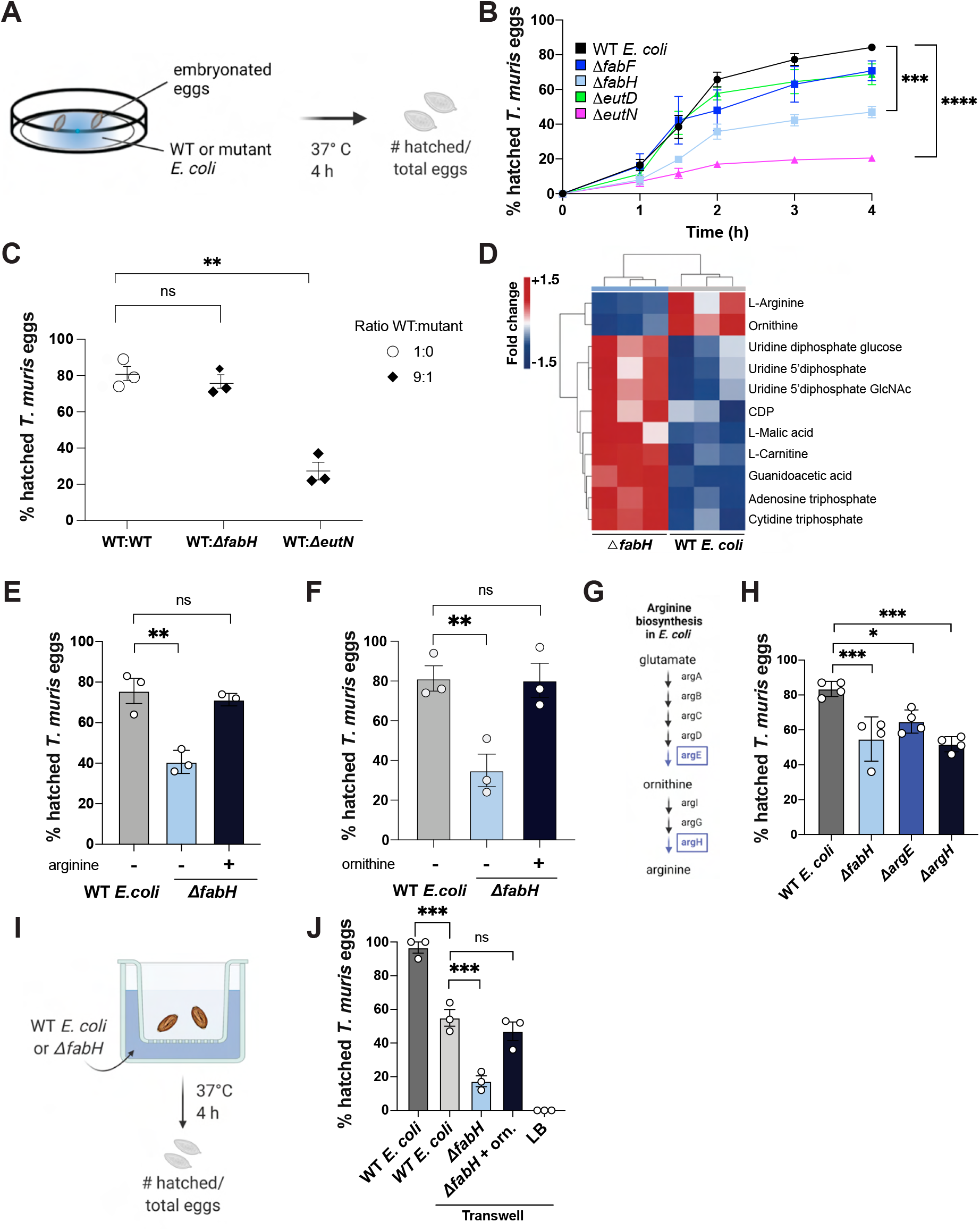
*E. coli* mutants with altered metabolic byproducts delay *C. elegans* fertility and inhibit *T. muris* egg hatching. **(A)** Experimental approach for *in vitro* hatching assay. **(B)** Percent *in vitro* hatched *T. muris* eggs after incubation with overnight cultures of four *E. coli* mutants identified from the *C. elegans* screen or WT *E. coli*. Hatched eggs were checked using light microscopy at timepoints indicated (n = 4); h = hours. **(C)** Hatching rates for mixtures of WT *E. coli* and WT *E. coli* or either Δ*fabH* or Δ*eutN* overnight cultures in the ratios indicated. Total concentration of each mixture was maintained at 1x 10^8^ CFU/mL (n = 3). **(D)** Hierarchical clustering of metabolites that were significantly (p < 0.05, Student’s t-test) downregulated (blue) or upregulated (red) in *ΔfabH* relative to WT. **(E, F)** Percent *in vitro* hatched *T. muris* eggs elicited by Δ*fabH* and cultures supplemented with (E) 500µM arginine or (F) 100µM ornithine where indicated (n=3). **(G)** Arginine biosynthesis pathway in *E. coli*. **(H)** Hatching rates elicited by Δ*argE* and Δ*argH* (n = 4). **(I)** Experimental approach for hatching assay in a transwell system. **(J)** Hatching quantified for eggs contained within a transwell chamber separated from bacterial culture in the outer chamber by a membrane with 0.4µm pores. Samples supplemented with ornithine as in (H) are indicated (n = 3). (B) One-way analysis of variance (ANOVA) of area under curve compared to WT. (C, E-F, H, J) One-way analysis of variance (ANOVA) with Dunnett’s post-test compared to WT. For all panels, measurements were taken from distinct samples. Graphs show means and SEM. * p < 0.05, ** p < 0.01, *** p < 0.001, **** p < 0.0001. ns = not significant.

To determine whether reduced egg hatching is due to the production of a toxic factor, we performed the hatching assay utilizing WT *E. coli* mixed with each mutant (Fig. 2C). As little as 10% *ΔeutN* in the mixture significantly inhibited the ability of WT *E. coli* to mediate hatching, whereas the presence of *ΔfabH* had no effect. Thus, the inhibitory effect of *ΔeutN* is dominant over WT *E. coli*, implicating the production of a toxic factor by this mutant, while the reduced hatching rate of embryonated eggs incubated with *ΔfabH* is likely due to deficiency of a pro-hatching factor.

Hybrid metabolomics revealed an unexpectedly low relative level of arginine and ornithine in *ΔfabH* cell lysates compared to WT *E. coli* (Fig. 2D). Although interactions between the fatty acid and arginine biosynthesis pathways of *E. coli* have not been previously described, *fabH* deficiency would cause a buildup of its substrate malonyl-ACP. A possible consequence of this buildup is compensatory decreased cycling of acetyl-CoA, particularly through the TCA cycle, and consequential slowing of the urea cycle and arginine biosynthesis. Supplementation of *ΔfabH* cultures with arginine or ornithine successfully rescued *T. muris* egg hatching to levels similar to WT *E. coli* (Fig. 2, E-F). Similarly, supplementation of *ΔfabH* cultures with arginine or ornithine significantly increased *C. elegans* fertility (Fig. S2, D-E). Consistent with a role for the arginine biosynthesis pathway, we found that *Trichuris* egg hatching was diminished in the presence of *ΔargE* or *ΔargH E. coli*, mutants defective in ornithine and arginine synthesis, respectively (Fig. 2, G-H).

Although optimal *T. muris* egg hatching *in vitro* requires physical contact with *E. coli* (Hayes et al., 2010; Koyama, 2016), a reduced amount of hatching can still occur in a transwell system that separates bacteria and eggs into different chambers (Koyama, 2016), suggesting that both contact-dependent and independent factors contribute to the process. Similar to previous reports, eggs hatched at approximately 50% their maximum rate when incubated separately from WT *E. coli* in a transwell system (Fig. 2, I-J). Hatching was further reduced when *ΔfabH* was used as the bacterium. Supplementation of ornithine restored hatching in the presence of *ΔfabH* to levels similar to those elicited by WT *E. coli* in the transwell system. Altogether, these findings indicate that nematode pathways regulated by ornithine and arginine contribute to contact-independent *E. coli*-mediated hatching of *T. muris* embryonated eggs.

### Toxic intermediates of the *E. coli* ethanolamine utilization pathway inhibit *T. muris* egg hatching

To understand why *T. muris* eggs incubated with *ΔeutN E. coli* displayed the greatest reduction in hatching, we first determined whether disrupting other parts of the ethanolamine utilization pathway (Fig. 3A) led to similar outcomes. *E. coli* mutants lacking enzymes in the pathway—*ΔeutB*, *ΔeutC*, *ΔeutD*, *ΔeutE*, and *ΔeutG*—facilitated egg hatching at rates similar to WT *E. coli* (Fig. 3B). In contrast to these *eut* operon genes, *eutN* encodes a subunit of a proteinaceous shell that enables the efficient colocalization of the *eut* enzymes with their cofactors and substrates within a microcompartment and prevents diffusion of pathway intermediates (Yeates et al., 2010). It is possible that disruption of the microcompartment architecture allows the release of a soluble toxic intermediate. Therefore, we tested whether transferring the supernatant from *ΔeutN* culture was sufficient to inhibit the ability of WT *E. coli* to mediate hatching (Fig. 3C). Although transferring culture supernatant from *ΔeutN* grown in isolation had no effect, supernatant isolated from *ΔeutN* that had been incubated with eggs (*ΔeutN^egg^*) reduced hatching mediated by WT *E. coli* (Fig. 3D). This suggests that *ΔeutN* produces a soluble toxic factor in the presence of *T. muris* embryonated eggs.

**Fig. 3.**
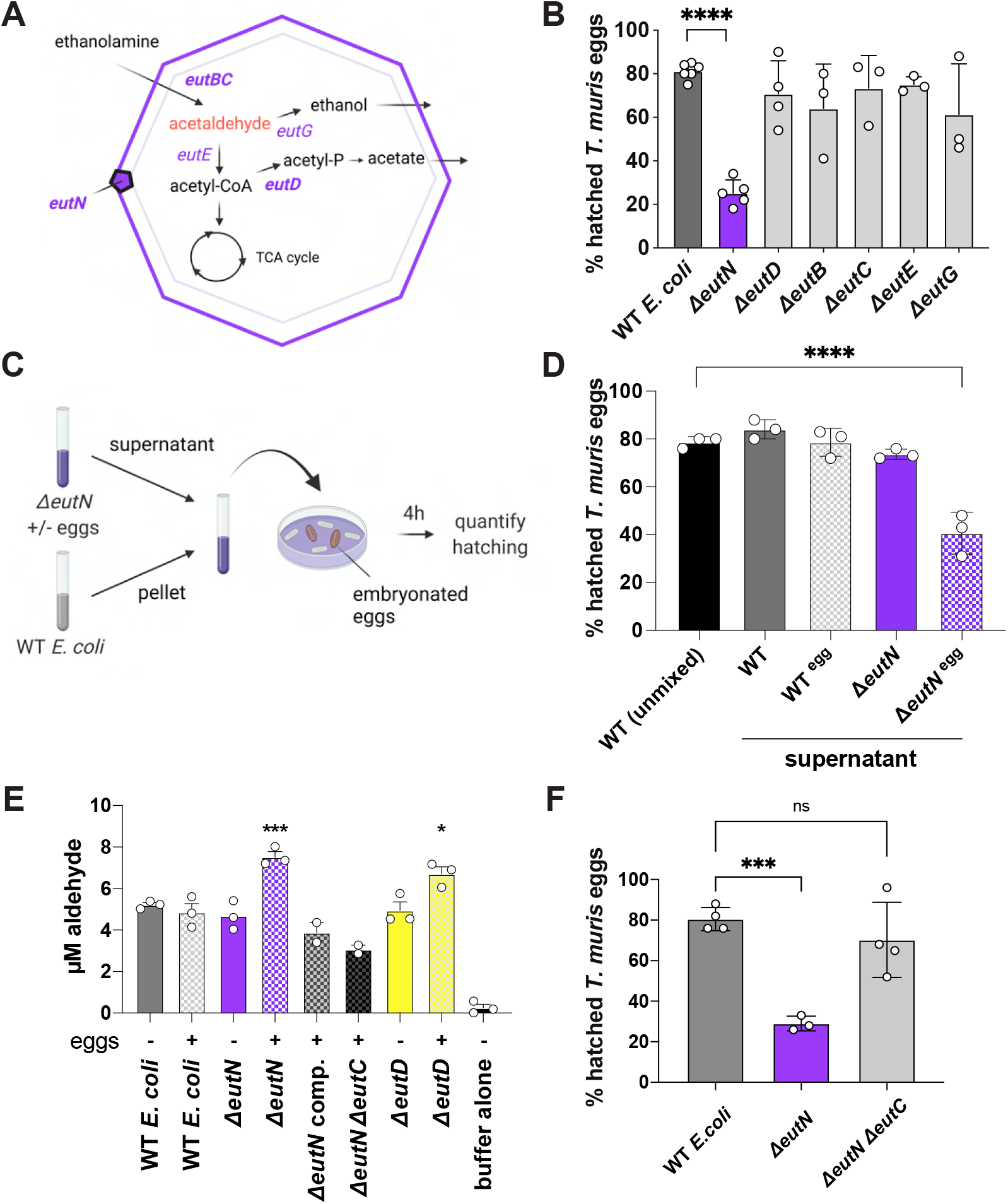
Toxic intermediates of the *E. coli* ethanolamine utilization pathway inhibit *T. muris* egg hatching. **(A)** Ethanolamine utilization pathway in *E. coli*. Modified from (Kaval and Garsin, 2018). **(B)** Hatching rates for *eut* mutants in the Keio library and WT controls (n = 3-6). **(C)** Experimental approach to assess the toxicity of soluble factors produced by *E. coli* Δ*eutN*. Filtered supernatant from Δ*eutN E. coli* or controls grown with or without *T. muris* eggs for four hours were transferred to a dish containing WT *E. coli* and *T. muris* eggs. The ability of WT *E. coli* to mediate egg hatching in the presence of supernatant was examined four hours later. **(D)** Quantification of hatching rates for mixtures containing WT *E. coli* cells and the indicated filtered supernatants (n = 3). **(E)** Aldehyde concentrations in supernatants of indicated *E. coli* strains incubated with or without *T. muris* eggs; Δ*eutN* comp. = Δ*eutN* strain where *eutN* has been complemented on a plasmid. **(F)** Hatching rates for Δ*eutN*Δ*eutC* double mutant (n = 4). (B, D-F) Measurements were taken from distinct samples. The average per independent experiment for at least three technical replicates shown. Graphs show means and SEM. One-way analysis of variance (ANOVA) with Dunnett’s post-test compared to WT. * p < 0.05, *** p < 0.001, **** p < 0.0001. ns = not significant.

Acetaldehyde is a known intermediate of the ethanolamine utilization pathway that can readily form irreversible adducts with proteins, changing their conformation and function (Donohue et al., 1983; Sorrell and Tuma, 1987). Additionally, deletion of *eutN* in *Salmonella* resulted in the greatest increase in acetaldehyde volatility among all *eut* operon mutants during the first hours of growth (Penrod and Roth, 2006). Using a fluorogenic hydrazone transfer (DarkZone) system to label aldehydes (Yuen et al., 2016), we detected elevated levels of aldehydes in the culture supernatant of *ΔeutN^egg^*, but not in the culture supernatant of the complemented *ΔeutN* strain or when the mutant was grown in the absence of eggs (Fig. 3E and S3A). Aldehyde levels were also elevated in *ΔeutD* (Fig. 3E), the other *E. coli* mutant of this pathway identified by the *C. elegans* screen, although the increase following egg incubation was not as striking as for *ΔeutN*, potentially explaining why deletion of *eutD* did not inhibit hatching (Fig. 2B).

The eut operon transcriptional regulator in *E. coli*, *eutR*, is sensitive to the chemical environment (Penrod and Roth, 2006). We hypothesized that *eutR* expression may be altered in the presence of *T. muris* eggs, thus explaining why aldehyde levels are increased when bacteria are co-incubated with eggs. We found that *eutR* expression in WT *E. coli* and *ΔeutN* significantly increased following egg incubation (Fig. S3B). As a control, increased *eutR* expression was detected from WT *E. coli* grown in minimal medium supplemented with ethanolamine, consistent with previous studies (Penrod and Roth, 2006; Rowley et al., 2020).

Next, we deleted the rate limiting enzyme of the ethanolamine utilization pathway, *eutC*, in the *ΔeutN* background (*ΔeutNΔeutC)* to prevent the formation of toxic intermediates (Penrod and Roth, 2006; Roof and Roth, 1989). We found that aldehyde levels of *ΔeutNΔeutC^egg^* were similar to that of WT *E. coli* and that *T. muris* egg hatching was completely restored when the assay was performed with this double mutant (Fig. 3E-F). We also found that pre-treatment of *ΔeutN^egg^* supernatant with an aldehyde scavenger N-acetyl cysteine completely abrogated its effects on WT *E. coli* and restored hatching (S3C-D). Additionally, we found that supplementation of acetaldehyde alone was sufficient to inhibit hatching elicited by WT *E. coli* in a concentration-dependent manner and confirmed that bacteria viability was unaffected (Fig. S3, E-F).

### *T. muris* reared in mice colonized with *E. coli* mutants exhibit defects in worm morphology and fertility

*In vitro T. muris* egg hatching serves as a sensitive assay to investigate nematode-bacteria interactions, but the role of intestinal bacteria in subsequent steps of *Trichuris* development remains obscure because they are difficult to recreate outside the mammalian host. Given that the *C. elegans* screen was focused on development of the reproductive tract, we established a bacterial monocolonization mouse model to investigate the role of bacterial *fabH*, *eutN*, and *eutD* in *T. muris* development and reproduction (Fig. 4A). After we verified that each *E. coli* mutant colonized germ-free mice to levels similar to WT *E. coli* (Fig. S4A), monocolonized mice were gavaged twice with 100 eggs and sacrificed two weeks after the second gavage. *E. coli* colonization and cecum length were confirmed to be similar across groups on the day of sacrifice (42 dpi) (Fig. S4, A-B).

**Fig. 4.**
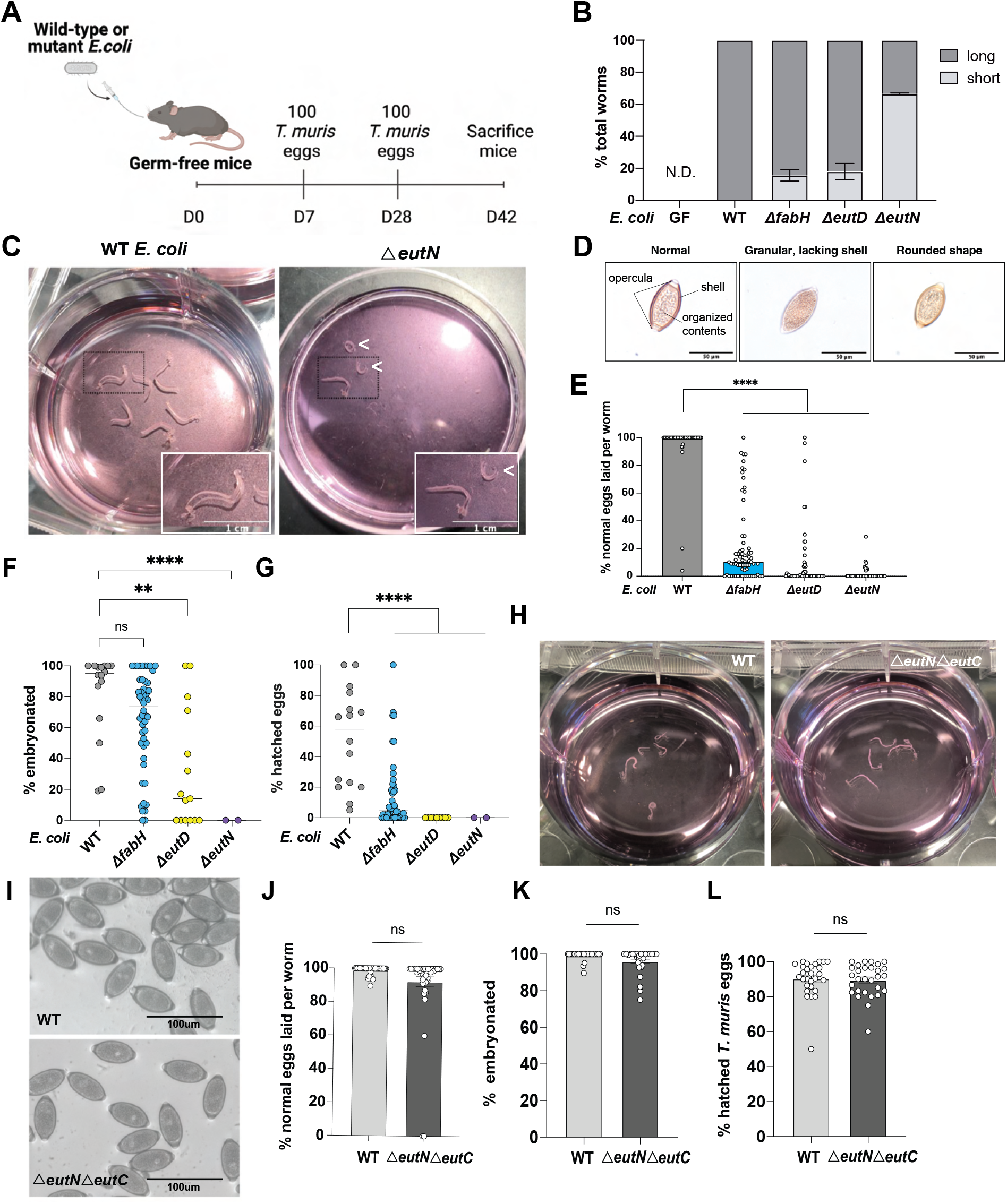
*T. muris* reared in mice colonized with *E. coli* mutants exhibit defects in worm morphology and fertility. **(A)** Experimental approach for *T. muris* infection of mice monocolonized with WT or mutant *E. coli*. **(B)** Proportion of short and long worms harvested per mouse colonized by indicated *E. coli* strains; N.D. = not detected (n = 6-9 mice per group). **(C)** Representative images of worms harvested from mice monocolonized with WT or *ΔeutN E. coli*. A representative short worm (indicated by arrowhead) next to a long worm is shown in the inset for *ΔeutN E. coli*. **(D)** Representative images of eggs laid by worms from (B). Worms were incubated in RPMI medium overnight to trigger egg laying. **(E)** Proportion of total eggs laid per worm with normal morphology. Circles represent individual worms (WT, n = 46; *ΔfabH*, n = 68; *ΔeutD*, n = 58; *ΔeutN*, n = 50). **(F)** Percentage of embryonated eggs following 8-week incubation. **(G)** Percentage of eggs in (F) that hatched after incubation with WT *E. coli* for 4 hours at 37°C. **(H)** Representative photos of worms harvested from WT and *ΔeutNΔeutC* monocolonized C57BL/6 mice. **(I)** Representative images of eggs laid by worms harvested from monocolonized mice groups indicated**. (J)** Proportion of total eggs laid per worm with normal morphology**. (K)** Percentage of embryonated eggs following 8-week incubation. **(L)** Percentage of eggs in (K) that hatched after incubation with WT *E. coli* for 4 hours at 37°C. (B, E, F-G) Graphs show (B) means and SEM or (E-G) medians. One-way analysis of variance (ANOVA) with Dunnett’s post-test compared to WT. (J-L) Graphs show means and SEM. Two-tailed, unpaired t-test. For all panels, measurements were taken from distinct samples. Data shown are pooled from two independent experiments. ** p < 0.01, **** p < 0.0001. ns = not significant.

We were able to recover adult worms from the cecum of mice monocolonized with WT *E. coli* but not germ-free mice (Fig. S4C), consistent with previous observations (White et al., 2018), indicating that our assay allows investigation of mechanisms by which *E. coli* contributes to *T. muris* development in an intact host. Our experimental setup in which mice receive high doses of embryonated eggs was designed to overcome the potential reduced hatching efficiency that may occur in mice lacking a complex microbiota. Therefore, as expected, we did not detect any differences in worm burden when comparing mice colonized with WT *E. coli* versus *ΔfabH*, *ΔeutN*, and *ΔeutD* strains (Fig. S4C). However, many of the *T. muris* recovered from mice colonized with these mutants displayed a striking deformity characterized by shortened length (Fig. 4, B-C; indicated by arrowheads). Although worms from WT *E. coli*-colonized mice were exclusively long in length, mice colonized with *ΔfabH*, *ΔeutN*, and *ΔeutD E. coli* produced a mixture of both long and short length worms, defined as < 1cm. The defect was most severe in mice monocolonized with *ΔeutN* from which most worms recovered were short. To investigate if the shortened length of *T. muris* resulted from mutant *E. coli* colonization affecting the host, we examined T cell subsets that have been previously established as key determinants of host susceptibility (Bancroft et al., 1994; D’Elia et al., 2009). Flow cytometric comparison of mesenteric lymph node T cells from mice colonized with WT or *ΔeutN E. coli* revealed no significant differences in T helper type 1 (Th1), T helper type 2 (Th2), regulatory T (T reg) cells, or IFN-γ producing T cells (Fig. S4D-G).

The average number of freshly laid eggs per worm was not significantly different between the conditions or when comparing long versus short worms (Figure S4H). However, in stark contrast to worms from WT *E. coli*-colonized mice, which laid eggs of the typical ovoid structure, worms harvested from mice colonized with *ΔfabH*, *ΔeutD*, and *ΔeutN E. coli* laid more eggs that displayed an aberrant rounded morphology, and in some cases lacked opercula and a distinct shell, and the inner contents were disorganized (Fig. 4, D-E). These structurally aberrant eggs took longer to reach embryonated morphology, and even after a prolonged incubation period, significantly fewer eggs appeared embryonated in the *ΔeutN* and *ΔeutD* groups (Fig. 4F). Eggs laid by worms from all three mutant-colonized groups of mice had significantly lower hatching rates compared to eggs laid by worms from WT *E. coli*-colonized mice when tested for bacteria-mediated hatching *in vitro* (Fig. 4G). These hatching defects persisted even when eggs were allowed to embryonate twice the embryonation time published in the literature for *T. muris* (Fig. S4I), suggesting disrupted and not delayed development. In line with our *in vitro* data (Fig. 3F), monocolonization of mice with *ΔeutNΔeutC* abrogated the length and egg fitness defects of *T. muris* observed in *ΔeutN-*colonized mice (Fig. 4, H-L). A similar rescue was seen when mice were monocolonized with the *ΔeutN* complemented strain (Fig S4, J-L). These findings indicating that bacterial deficiencies decrease *T. muris* worm size and egg fitness is consistent with recent evidence showing that another parasitic nematode *Heligmosomoides polygyrus*, lays fewer eggs and exhibits similar length defects when reared in a germ-free host (Rausch et al., 2018).

To gain insight into the cause of morphologic and reproductive aberrancies in *T. muris*, we performed RNA-Seq analysis of individual female worms (Fig S4M, Table S2). The transcriptomes of worms harvested from WT *E. coli*-colonized mice clustered separately from those of short and long worms harvested from mutant-colonized mice. The latter did not segregate further according to length, representing no detectable significant differences between short and long worms particularly in developmental genes. Analysis of the top 50 most differentially detected genes between the WT- and mutant-associated worms (Table S3) included genes related to *C. elegans* genes *egg-3*, *egg-4*, *egg-5*, *alg-3*, and *alg-4* all of which are expressed in the germ line of *C. elegans* and whose biological functions include roles in oogenesis, spermatogenesis, fertilization, the egg-to-oocyte transition, and egg shell deposition (Boeck et al., 2016; Cheng et al., 2009; Conine et al., 2010; Maruyama et al., 2007; Parry et al., 2009). Crosschecking our 50 genes with published lifestage- and sex-specific transcriptomics of *T. muris* (Foth et al., 2014), all genes found to be highly expressed in the female worms from mutant-colonized mice were previously detected as upregulated in adults (vs. larvae), providing additional evidence that the short worms are not disrupted in somatic development. Additionally, these same genes were also previously detected as more highly expressed in *T. muris* males compared to females (Foth et al., 2014). Conversely, all but one of the genes that were downregulated in the worms from mutant-colonized mice were found to be upregulated in *T. muris* females. These discordances provide evidence of sexual development dysregulation in worms harvested from mutant-colonized mice.

In conclusion, these results provide evidence for a previously undescribed role for byproducts of microbial metabolism on nematode development, a role that appears evolutionarily conserved. Despite lacking many genes specific to parasitism, our findings support the utility of *C. elegans* as a model for bacteria-nematode interactions because of its high sensitivity to its bacterial milieu. In addition to identifying two critical pathways in bacteria, fatty acid biosynthesis and ethanolamine utilization, our findings describe a novel pipeline that leverages genetic tools available in *C. elegans* to discover new helminth biology. An important future direction would be to determine the role of microbial byproducts in the setting of a complex microbiota. For instance, microbiota rich in bacteria with highly active arginine biosynthesis or another pathway favorable for *T. muris* reproduction may confer greater susceptibility to infection. Identifying the microbes and their products that affect different stages of the helminth life cycle would allow predictions and testable hypotheses regarding the impact of interindividual differences in microbiota composition. We propose that *C. elegans* can be used as a system for investigating the effects of specific microbial genes and pathways in supporting the parasitic nematode life cycle and aid in further understanding the transkingdom interactions that sustain human disease.

## Acknowledgements

We wish to thank D. Jones, R. Rose, and L. Ash from the NYULH Metabolomics Laboratory (SCR_017935) and C. Yun, S. Cook, and D. Kahler from the High Throughput Biology Core (now Microscopy Laboratory, NIH S10 RR023704) for their help in acquiring and analyzing the data presented. Both Core Laboratories are funded partially by NIH/NCI P30CA016087. Some strains were built using strains provided by the CGC which is funded by the NIH Office Research Infrastructure Programs (P40 OD010440), and we thank WormBase. We also thank H. Darwin and M. Pacold for advice on microbial metabolism; S. Marion, C. Linton, M. Xu, A. Niranjan, D. Stokes, M. Lopez-Redondo, I. Irnov, V. Torres, S. Dyzenhaus, M. Hui, J. Richards (NYU), and E. Tait-Wojno (Univ. of Washington) for technical assistance and reagents; members of the Hubbard Lab, Cadwell Lab, and NYU Medical Scientist Training Program for constructive comments and technical assistance; and M. Alva, J. Carrasquillo, and D. Basnight for assistance with gnotobiotics.

## Funding

NIH grant DK093668 (K.C.)

NIH grant AI121244 (K.C.)

NIH grant HL123340 (K.C.)

NIH grant AI130945 (K.C.)

NIH grant AI140754 (K.C.)

NIH grant DK124336 (K.C.)

NIH grant R01GM130152 (E.J.A.H.)

NIH grant R35GM134876 (E.J.A.H.)

NIH grant R01GM035769 (J.G.B.)

NIH grant 5T32AI100853 (M.V.)

NIH grant T32GM136573 (M.V.)

NIH grant T32GM007308 (M.V.)

Faculty Scholar grant from the Howard Hughes Medical Institute (K.C.)

Crohn’s & Colitis Foundation (K.C.)

Kenneth Rainin Foundation (K.C.)

Judith & Stewart Colton Center of Autoimmunity (K.C.)

US National Cancer Institute (CA217809) (E.T.K.)

K.C. is a Burroughs Wellcome Fund Investigator in the Pathogenesis of Infectious Diseases.

## Author contributions

M.V., R.D., E.J.A.H, and K.C. conceived the study and designed the experiments. M.V., R.D., and J.B. performed, analyzed, and interpreted all the experiments for *T. muris* and *C. elegans*. D.J.L. performed the bacterial cloning. H.S.P and E.T.K provided DarkZone and technical advice. J.C.D. provided technical assistance and advice for RNA sequencing and analysis. K.C., E.J.A.H, and J.G.B oversaw analysis and interpretation of all experiments described. M.V., R.D., E.J.A.H., and K.C. wrote the manuscript with inputs from all authors.

## Data and materials availability

All data needed to evaluate the conclusions in this study are available in the main text and the supplementary materials or available from the corresponding authors upon request.

## Supplementary Materials

Methods

Figs. S1 to S4

Tables S1-S7

## Methods

### Keio Library growth and culture

The frozen Keio collection was stamped out onto 150mm LB kanamycin (30µg/ml) plates and grown overnight at 37^°^C. For *C. elegans* experiments, individual colonies were inoculated into 700 µl of LB-Kan (30µg/ml) in 96 deep well plates using a compatible pin replicator. The plates were sealed with pure link air porous tape (Invitrogen cat no. 12262010) and incubated for 16-18 hours at 37^°^C, shaking at 250 rpm. Cultures were diluted 10X in S-media in 96 well format and OD at 600 nm was recorded for future reference to ensure that candidate positive mutants grew to an OD similar to the GC1547 wild-type control wells in the same plate. (This was done since poor bacterial growth can reduce the percentage of gravid *C. elegans*; all 315 candidates from the primary screen passed this control). Bacterial cultures were pelleted by centrifugation at 2000 rpm for 30 minutes. Supernatant was discarded and the pellet was resuspended in 100µl of S-media. 40µl of the culture was transferred into two separate plates (two technical replicates; black-walled, clear bottom 96-well microplates; Corning Cat. No. 3904) using a multi-channel pipette. The GC1547 control wild-type strain was a kanamycin resistant transformant of the Keio library starting strain BW25113, carrying the pET28a plasmid. For *T. muris in vitro* hatching assays, *E. coli* was grown overnight for 12-14 hours in Luria-Bertani broth with shaking at 225 rpm at 37 °C.

### Primary screen

*C. elegans* GC1474 (Table S1; derived by outcrossing GC1373 (Roy et al., 2018) of the genotype *glp-1(e2141); end-1::gfp; myo-2p::mCherry*) was maintained on OP50 *E. coli* on NGM plates at 20°C and synchronized in the L1 larval stage using hypochlorite treatment followed by overnight hatching in egg buffer (Shaham, 2006). 40-50 L1 larvae were seeded into 96-well plates in technical duplicate with 40µl of *E. coli* mutant cultures and incubated at 20^°^C with mild shaking in a humidified chamber for 62-65 hours. At this timepoint, control worms were fully gravid adults; control GC1547 bacteria were included on each plate in wells void of library bacteria. Prior to imaging, 40µl of 2mM levamisole was added to each well to immobilize the worms. The plates were then sealed with aluminum foil (Corning Cat. No. 6570) and imaged to detect both green (embryos) and red (pharynx) fluorescent markers using arrayscan (Thermo Scientific VTI). Images were captured at 2.5X magnification 2×2 binning in a 16 mm^2^ field and digitally archived. Images captured on the red and green channels were merged in ImageJ using macros written to automate the process. Each adult animal was scored visually as “with embryos” (green embryos visible in the uterus) or “without embryos” (no green embryos visible in the uterus), and the number of worms without embryos and the total number of worms (excluding any that were arrested or any that crossed the edge of the well). Wells in which many animals had not reached adulthood were marked and were not considered further in our analysis. Ten such bacterial mutants grew well by OD but did not support worm development.

Additional culling of potential hits was performed. A variety of technical factors eliminated fewer than 10% of the total: 54 library wells were devoid of bacteria as per the plate map; 58 had a gene name in the library but the bacteria did not grow (either on solid or liquid culture); 41 were missed due to human error while scoring (noted as largely fertile but not counted individually); 5 had low n; 4 were not included due to reasons classified as ‘other’ (e.g., poor image) and 10 wells had poor worm growth in both replicates.

Worm counts (with/without embryos) were exported into ActivityBase (IDBS, https://www.idbs.com; a Scientific informatics software platform) and the robust Z score (a statistical measure that takes into account the median plate “without embryos” percent) was calculated for every well to minimize problems associated with plate to plate variation. Each well was analyzed with respect to percent worms without embryos, consistency (across technical replicates) and robust Z score. A cut-off of robust Z ≥ 1.5 was selected as “Z positive” (≤ 10% of the wells per plate scored above this threshold). We also calculated the penetrance of worms without embryos for pooled technical replicates.

Criteria for selection in the primary screen weighed the robust Z and penetrance, with special consideration for cases where only one replicate was available. We included those wells for which only one technical replicate was available provided that well made the RobZ cut off (≥ 1.5) and/or percent non-gravid worms as further described below. Reasons for single-replicate included low n (62 wells), human error (49 wells), worm arrest (31 wells), poor image quality (22 wells) and 1 well marked as ‘other’. Considering Z score, penetrance and replicates, the candidates were further classified as follows: High, A: both replicates Z positive and average penetrance of non-gravid worms > 20%, or B: one replicate available and it is Z positive and > 20%, or C: both replicates available but only one is Z positive and non-gravid ≥ 30%. Medium, A: both replicates Z positive and pooled non-gravid percent 13-20%, or B: only one replicate available which is Z positive and 13-20% penetrance of worms without embryos, or C: both replicates available but only one is Z positive with pooled non-gravid penetrance between 20-30%. Low: both replicates available, but only one is Z positive and non-gravid percent between 12.5-20%. This analysis yielded 315 primary screen candidates.

We then plotted the percent non-gravid in descending order for every well per plate and noted the inflection point (see (Roy et al., 2018)). We placed a subset of the mutants that passed the criteria above onto these plots and observed a strong correlation: wells with larger Z scores all appeared above the inflection point.

### Secondary screen: reproducibility

The 315 candidates from the primary screen were re-selected from the original Keio library and gridded onto secondary screen plates. Two plates included only high priority candidates or a combination of randomly selected high, medium and low priority candidates to help assess whether there was a correlation between penetrance and reproducibility. No such correlation emerged, but these plates were considered as additional replicates in the analysis going forward (hence 6 plates) though 315 mutants. Control bacteria were included in all secondary screen plates. Three biological replicates were performed in technical triplicate and scored and analyzed as described above. Technical replicate non-gravid/gravid counts were pooled and analyzed in pairwise comparison with the control on the same plates using Fisher’s exact test. Candidates with p-values ≤ 0.05 in at least 2 out of 3 biological replicates were chosen for further analysis. This analysis yielded 10 candidates.

### PCR validation of selected Keio mutants (completed for all ten hits)

We amplified and sequenced PCR fragments ∼150 bp upstream and downstream of the predicted gene from each mutant and the control. Nested primers were used to sequence the PCR fragment and confirm the presence of the FRT and kanamycin cassette in the mutant DNA (Tables S3, S5).

### P1 transduction

Mutant alleles comprising a kanamycin-resistance marker within a gene deletion were introduced into BW25113 by P1 transduction (Thomason et al., 2007) from the Keio collection (Baba et al., 2006). The kanamycin-resistance cassette was subsequently excised to generate a markerless in-frame gene deletion by transformation with the temperature-sensitive plasmid pCP20 (Cherepanov and Wackernagel, 1995), which encodes FLP recombinase, and growth at 30°C. Finally, pCP20 was eliminated from the cells by growth at 42°C.

### Complementation

For *fabH* and *eutN*, the bacterial gene coding regions plus 20 bp upstream were inserted into pBR322 downstream of a plasmid promoter by Gibson Assembly. The wild-type genomic DNA was amplified and assembled with a PCR fragment obtained from the pBR322 vector (specific primers are listed in Table S5; the fragment excluded part of the Amp^R^ gene between *Eco*RI and *Pst*I restriction site). PCR products were purified using gel extraction kit (QIAEX II cat no. 20021) and 100 ng of vector was used in 1:1 molar reaction with the genomic fragment. The assembly reaction was then transformed into high efficiency DH5alpha cells (NEB C2987I). For *eutD*, the protein-coding region was amplified by PCR and cloned between the NdeI and XbaI sites downstream of the *lac*UV5 promoter of plasmid pPlacRppH6 so as to replace the *rppH* coding region (Deana et al., 2008). Plasmid preparations (Qiagen mini prep kit Cat no.27106) were sequenced using plasmid-specific primers (Table S5). Plasmids were confirmed by DNA sequence and transformed into the respective mutant (so as to create strains GC1558, GC1559, and GC1650 for *eutN* rescue, *fabH* rescue, and *eutD* rescue, respectively) and tested for reduced penetrance of the *C. elegans* non-gravid phenotype upon feeding.

### Developmental timing and germline analysis

#### Somatic development assay

*C. elegans* were maintained on OP50, synchronized by hypochlorite treatment and allowed to hatch at 20°C with shaking. 30-40 L1 larvae were seeded in 96 deep well plates with each well carrying a specific mutant clone. Worms were incubated at 20°C with shaking for 48 hours and monitored at 400X on a Zeiss Z1 Axio Imager for vulval morphology (Mok et al., 2015) at specific time points.

#### Pro phenotype assay

*glp-1(ar202)* animals were scored for the Pro phenotype upon feeding with mutant bacteria using the same strategy as for the primary screen. Worms were maintained at 15°C on OP50 synchronized and grown in 96 deep well format as above. Day 1 (∼72 hours post seeding) worms were fixed and DAPI stained and each gonad arm was scored for the Pro phenotype (Pepper et al., 2003) at 400X.

#### Fertility at Day 2

GC1474 worms were maintained at 20°C on OP50, synchronized as L1 larvae and grown in 96 deep well format as above. Worms were anesthetized and imaged as in the screen on Day 1 and Day 2 to determine changes in penetrance of fertile worms.

#### Progenitor zone nuclei counts at L4-Adult molt

Worms grown on GC1547 control, *eutN*, *eutD,* and *fabH* mutant bacteria were monitored for vulval development and isolated at the L4-adult molt. Worms were then stained with DAPI (Michaelson et al., 2010) and 0.5µm Z-stack images were collected and analyzed using ImageJ to determine the number of germ cells in the PZ. In approximately 30% of the worms, the transition zone was not clear. These were censored from the analysis.

#### Assay for dependence on DAF-7/TGFß and DAF-2 IIS pathways

*C. elegans* strains GC1474, GC1545 and GC1250 (Table S2) were hypochlorite-synchronized and hatched overnight at 20°C. L1 larvae were seeded into 96 deep well plates containing specific bacterial mutants and scored for the presence of embryos in adults on Day 1 on Zeiss compound microscope at 100X.

### Parasite maintenance

Stock eggs of *T. muris* E strain (Ramanan et al., 2016) were maintained in the NOD.Cg-*Prkdc^scid^*/J (Jax) mouse strain in a specific pathogen free (SPF) facility and propagated as previously described (Antignano et al., 2011). Each egg batch was confirmed to hatch at ≥ 80% *in vitro* using method below and WT *E. coli* before use in subsequent experiments.

#### In vitro hatching of T. muris eggs

*T. muris* eggs were hatched *in vitro* by mixing 25µL of embryonated eggs at a concentration of 1 egg/1µL suspended in sterile water with 10µL of *E. coli* overnight culture and 15µL sterile LB in individual wells of a 48 well plate. Plates were incubated at 37°C and checked every hour for a total of four hours on the Zeiss Primovert microscope to enumerate hatched eggs. Rates describe hatching after four hours of incubation unless otherwise indicated. Experiments utilizing transwell inserts (Millicell) were performed as previously described (Koyama, 2016). For experiments where cell-free supernatant was used, supernatant and cells were isolated by centrifugation and filtration through a 0.22µm syringe filter or after wash with autoclaved ddH20, respectively. Incubation with embryonated eggs was performed by mixing *T. muris* eggs at a concentration of 15 eggs/2 mL culture and keeping at 37°C for four hours.

### Metabolomics

#### Sample preparation

Individual colonies (obtained after streaking frozen bacteria onto LB-kanamycin plates and incubating at 37°C) were inoculated in 30 ml of LB-kanamycin (30µg/ml) liquid in a 50ml falcon tube. Cultures were grown for 16-18hr at 37°C with 250 rpm shaking and then bacteria were pelleted by centrifugation at 1500 rpm for 20-30 minutes at 4°C. The supernatant was transferred to a fresh tube, filtered through 0.2µm filter (Corning (431229) and frozen at −80°C for future analysis. Pellets were resuspended in 7.5 ml of S-media (for ∼250ml of S-media the recipe was 240ml of S-basal, 2.5ml of 1M potassium citrate-final concentration of 10mM, 2.5ml of 1M trace metals for final concentration of 10mM, 750µl of 1M MgS0_4_ for final concentration of 3mM, 750µl of 1M CaCl_2_ for final concentration of 3mM, 72µl of 100mg/ml of kanamycin for final concentration of 30µg/ml). This solution was filter sterilized using a 500ml Nalgene filter (291-3320 Fisher). Post filter sterilization, 250µl of 5mg/ml cholesterol – final concentration of 5µg/ml and 2.5ml of 250µg/ml fungizone for final concentration of 2.5µg/ml (BP264550 Fisher).

The number of CFU/ml was estimated for each culture by plating dilutions (10µl of culture was mixed with 900µl of S-media to get 10^2^ or 1:100 dilution, which was then serially diluted to 1:10^5^ or 1:10^6^ on LB agar plates and counting colonies after growth at 37^°^C. The remaining culture was incubated at 20°C with mild shaking overnight to mimic *C. elegans* growth conditions. Based on CFU/ml measurements, 10^10^ cells were fast filtered (EZFIT Vacuum Manifold, Millipore EZFITLOW03 with Microfil V, Millipore MVHAWG124) and metabolite extracts were isolated as previously described (Irnov et al., 2017). Three independent biological samples were extracted on three independent days and subsequently analyzed by the NYU Langone Metabolomics Core Resource Library.

#### LC-MS/MS with the hybrid metabolomics method

Samples were subjected to an LCMS analysis to detect and quantify known peaks. A metabolite extraction was carried out on each sample based on a previously described method (Pacold et al., 2016). The LC column was a Millipore^TM^ ZIC-pHILIC (2.1 x150mm, 5μm) coupled to a Dionex Ultimate 3000^TM^ system and the column oven temperature was set to 25°C for the gradient elution. A flow rate of 100μL/min was used with the following buffers: A) 10mM ammonium carbonate in water, pH 9.0, and B) neat acetonitrile. The gradient profile was as follows; 80-20%B (0-30 min), 20-80%B (30-31 min), 80-80%B (31-42 min). Injection volume was set to 2μL for all analyses (42 min total run time per injection).

MS analyses were carried out by coupling the LC system to a Thermo Q Exactive HF^TM^ mass spectrometer operating in heated electrospray ionization mode (HESI). Method duration was 30 min with a polarity switching data-dependent Top 5 method for both positive and negative modes. Spray voltage for both positive and negative modes was 3.5kV and capillary temperature was set to 320°C with a sheath gas rate of 35, aux gas of 10, and max spray current of 100μA. The full MS scan for both polarities utilized 120,000 resolution with an AGC target of 3e6 and a maximum IT of 100ms, and the scan range was from 67-1000 *m*/*z*. Tandem MS spectra for both positive and negative mode used a resolution of 15,000, AGC target of 1e5, maximum IT of 50ms, isolation window of 0.4m/z, isolation offset of 0.1m/z, fixed first mass of 50m/z, and 3-way multiplexed normalized collision energies (nCE) of 10, 35, 80. The minimum AGC target was 1e4 with an intensity threshold of 2e5. All data were acquired in profile mode.

### Hybrid Metabolomics Data Processing

#### Relative quantification of metabolites

The resulting Thermo^TM^ RAW files were converted to mzXML format using ReAdW.exe version 4.3.1 to enable peak detection and quantification. The centroided data were searched using an in-house python script Mighty_skeleton version 0.0.2 and peak heights were extracted from the mzXML files based on a previously established library of metabolite retention times and accurate masses adapted from the Whitehead Institute (Chen et al., 2016), and verified with authentic standards and/or high resolution MS/MS spectral manually curated against the NIST14MS/MS (Simon-Manso et al., 2013) and METLIN (2017)(Smith et al., 2005) tandem mass spectral libraries. Metabolite peaks were extracted based on the theoretical *m*/*z* of the expected ion type e.g., [M+H]^+^, with a ±5 part-per-million (ppm) tolerance, and a ± 7.5 second peak apex retention time tolerance within an initial retention time search window of ± 0.5 min across the study samples. The resulting data matrix of metabolite intensities for all samples and blank controls was processed with an in-house statistical pipeline Metabolyze version 1.0 and final peak detection was calculated based on a signal to noise ratio (S/N) of 3X compared to blank controls, with a floor of 10,000 (arbitrary units). For samples where the peak intensity was lower than the blank threshold, metabolites were annotated as not detected, and the threshold value was imputed for any statistical comparisons to enable an estimate of the fold change as applicable. The resulting blank corrected data matrix was then used for all group-wise comparisons, and t-tests were performed with the Python SciPy (1.1.0) (Virtanen et al., 2020) library to test for differences and generate statistics for downstream analyses. Any metabolite with p-value < 0.05 was considered significantly regulated (up or down). Heatmaps were generated with hierarchical clustering performed on the imputed matrix values utilizing the R library pheatmap (1.0.12) (Kolde, 2015). Volcano plots were generated utilizing the R library, Manhattanly (0.2.0). In order to adjust for significant covariate effects (as applicable) in the experimental design the R package, DESeq2 (1.24.0) (Love et al., 2014) was used to test for significant differences. Data processing for this correction required the blank corrected matrix to be imputed with zeroes for non-detected values instead of the blank threshold to avoid false positives. This corrected matrix was then analyzed utilizing DESeq2 to calculate the adjusted p-value in the covariate model.

### Arginine and ornithine supplementation assay

*fabH* mutant *E. coli* or wild-type control was grown with 500µM or 100µM ornithine in LB-kanamycin overnight at 37°C with 250 rpm shaking. For *C. elegans* experiments, worms were imaged on Day 1 as described for the primary screen. For *T. muris* experiments, hatching assay was performed as described above.

### DarkZone labeling of aldehydes

Cell-free supernatants isolated from *E. coli* overnight cultures were incubated with 20µM DarkZone dye pre-dissolved in DMSO, 5mM 2,4 dimethoxyaniline catalyst (TCI America) pre-dissolved in DMSO, and buffer (100mM Tris pH 6.8, 150 mM NaCl) in a 96 well optical flat-bottom plate (Thermo-Fisher) in duplicate at 37°C for 1 hour. Adhesive plate seals were used to prevent evaporation of aldehydes. Fluorescence was measured using an EnVision 2 103 Multi-label Reader (PerkinElmer). Duplicate measurements were averaged.

### Quantitative PCR

Bacterial cells grown *in vitro* were pelleted and washed with RNA Protect (Qiagen) and PBS. For all samples, RNA was was isolated using the RNeasy micro kit with DNase treatment (Qiagen). Synthesis of cDNA was conducted with the High-Capacity cDNA Reverse Transcription Kit (Applied Biosystems). Samples were run without reverse transcriptase to control for genomic DNA contamination. Quantitative PCR was performed using SybrGreen (Roche) on a Roche480II Lightcycler using the following primers indicated in Table S5. Two technical replicates of each biological replicate were included for each gene target. Relative expression of the respective genes to *16S* rRNA expression was calculated using the ΔΔ*C*_T_ method.

### N-acetyl cysteine supplementation assay

Bacterial cells and cell-free supernatants of WT *E. coli* and *eutN* mutant *E. coli* were isolated as described above. A 100mg/mL stock concentration of N-acetyl cysteine (Millipore Sigma) was dissolved in MilliQ water by stirring at 70°C. The N-acetyl cysteine was then added to supernatants at the concentrations of 8µM, 10 µM, 30 µM, and 50 µM and incubated at room temperature for one hour. WT *E. coli* cells were then resuspended in the supernatants and incubated with *T. muris* eggs to perform the hatching assay as described above.

### Mice

#### Gnotobiotics

Previously described germ-free C57BL/6J mice (Martin et al., 2018) were maintained in flexible film isolators, and absence of faecal bacteria and fungi was confirmed by aerobic culture in brain heart infusion, sabaraud and nutrient broth (Sigma), and qPCR for bacterial 16S and eukaryotic 18S ribosomal RNA genes through sampling of stool from individual cages in each isolator on a monthly basis. Mice were transferred into individually ventilated Tecniplast ISOcages for infections to maintain sterility under positive air pressure. All animal studies were performed according to approved protocols and ethical guidelines established by the NYU Grossman School of Medicine Institutional Animal Care and Use Committee (IACUC) and Institutional Review Board.

#### Murine in vivo infections

Female mice were monocolonized at 6-8 weeks of age by oral gavage with 1 × 10^8^ colony forming units per mL (CFU/mL) of indicated *E. coli* strains. Overnight *E. coli* cultures were diluted 1:100 in Luria-Bertani broth followed by 2-3 hours of growth until 1 × 10^8^ CFU/mL was reached. Bacterial density was confirmed by dilution plating. Cultures were pelleted by centrifugation at 2437g for 10 minutes and washed once with sterile 1x PBS. Pellets were then resuspended in sterile 1x PBS and mice were inoculated by oral gavage with 1×10^8^ CFU in a volume of 100µL. Inoculum was verified using dilution plating of aliquots.

7 and 28 days later, mice were infected by oral gavage with ∼100 embryonated *T. muris* eggs. Individual worms were collected from cecal contents of all infected mice and washed in RPMI 1640 (Corning) supplemented with penicillin (100U/ml) and streptomycin (100μg/ml; Sigma). To evaluate egg laying, each worm was placed into individual wells of a 48 well plate with 200µL supplemented RPMI. Plates were then incubated at 37°C overnight in a closed tupperware (Sistema) lined with damp paper towels. The following day, eggs laid were enumerated using a Zeiss Primovert light microscope at 100X. Samples containing ∼1000 eggs per condition were mounted on glass slides with glycerol mounting medium (Sigma-Aldrich) and analyzed using a Zeiss Axio Observer at 400X with oil emersion for color images or EVOS FL Auto (Life Technologies) at 200X for black and white images. Images were processed using ImageJ.

### Flow Cytometry

Cells from mesenteric lymph nodes were isolated and stimulated as previously described (Ramanan et al., 2016). Stimulated cells were stained with anti-CD45 Pacific Blue, anti-CD3ε FITC, anti CD4 APC-Cy7, and anti-IFNγ AF700 from Biolegend and anti-CD19 PerCP Cy5.5 from eBioscience. Fixation and permeabilization buffers from Biolegend were used for intracellular cytokine staining, and a fixable live/dead stain from Biolegend was used to exclude dead cells. For nuclear staining, unstimulated cells were stained with anti-CD45 Pacific Blue, anti-CD3 FITC, anti-CD4 APC-Cy7, anti-FoxP3 PE-Cy7, and anti-Tbet APC from Biolegend, and anti-CD19 PerCP Cy5.5 and anti-GATA3 PE from eBioscience using the Foxp3 staining kit (eBioscience). Flow cytometric analysis was performed on a CytoFLEX analyzer (Beckman Coulter) and analyzed using FlowJo 10.0.8.

### RNA isolation from harvested *T. muris* and RNA-seq

*T. muris* harvested from mice ceca were immediately transferred to RPMI 1640 (Invitrogen) supplemented with penicillin (100U/ml) and streptomycin (100μg/ml; Sigma) and stored at −80°C. RNA from individual worms was isolated using the RNeasy micro kit with DNase treatment (Qiagen). RNA-seq library preparation was done at the NYU School of Medicine Genome Technology Core using an automated Trio enrichment library preparation protocol (NuGEN). Libraries were then sequenced on a NovaSeq 6000 (Illumina). All sequences have been deposited at NCBI under the BioProject PRJNA803354.

### RNA-seq analysis

Raw RNA-seq reads were trimmed using Trimmomatic v.036 (Bolger et al., 2014) and aligned to the reference *T. muris* genome (GenBank Assembly Accession: <u>G</u>CA_000612645.2) using the splice-aware STAR v2.7.3 (Dobin et al., 2013). The featureCounts command from Subread v1.6.3 (Liao et al., 2014) was used to generate counts for each gene based on how many aligned reads overlapped the exons of that gene. These counts were then normalized and used to test for differential expression using negative binomial generalized linear models implemented by DESeq2 v1.24.0 (Love et al., 2014).

### Functional gene annotation of RNA-Seq hits

Annotation of the 50 most differentially detected hits from the RNA-seq analysis described above was performed using WormBase (version WS283) searches and BLAST searches against the *C. elegans* reference genome using WormBase BLAST/BLAT. The E-value threshold was set at 1E+0 and the *C. elegans* Sequencing Consortium genome project database was used. *C. elegans* genes with the highest homology were noted and their functional descriptions derived using WormBase WormMine. Stage- and sex-specific expression of hits were assigned based on previously published data (Foth et al., 201cl4).

### Statistical analysis

Statistical tests and parameters used, including the definition of central value and the exact number (n) of mice or worms per group, are annotated in the corresponding figure legend. All analyses were performed with Graphpad Prism version 8.4.3 for Mac (GraphPad). The numbers of animals or biological replicates used herein were estimated on the basis of a power analysis with the following assumptions: the standard deviation will be roughly 20% of the mean; *p* values will be less than 0.05 when the null hypothesis is false; and the effect size (Cohen’s *d*) is between 1.0 and 2.0. We have also carefully chosen the indicated sample size on the basis of empirical evidence of what is necessary to interpret the data and statistical significance.

## Supplementary Figures

**Fig. S1.**
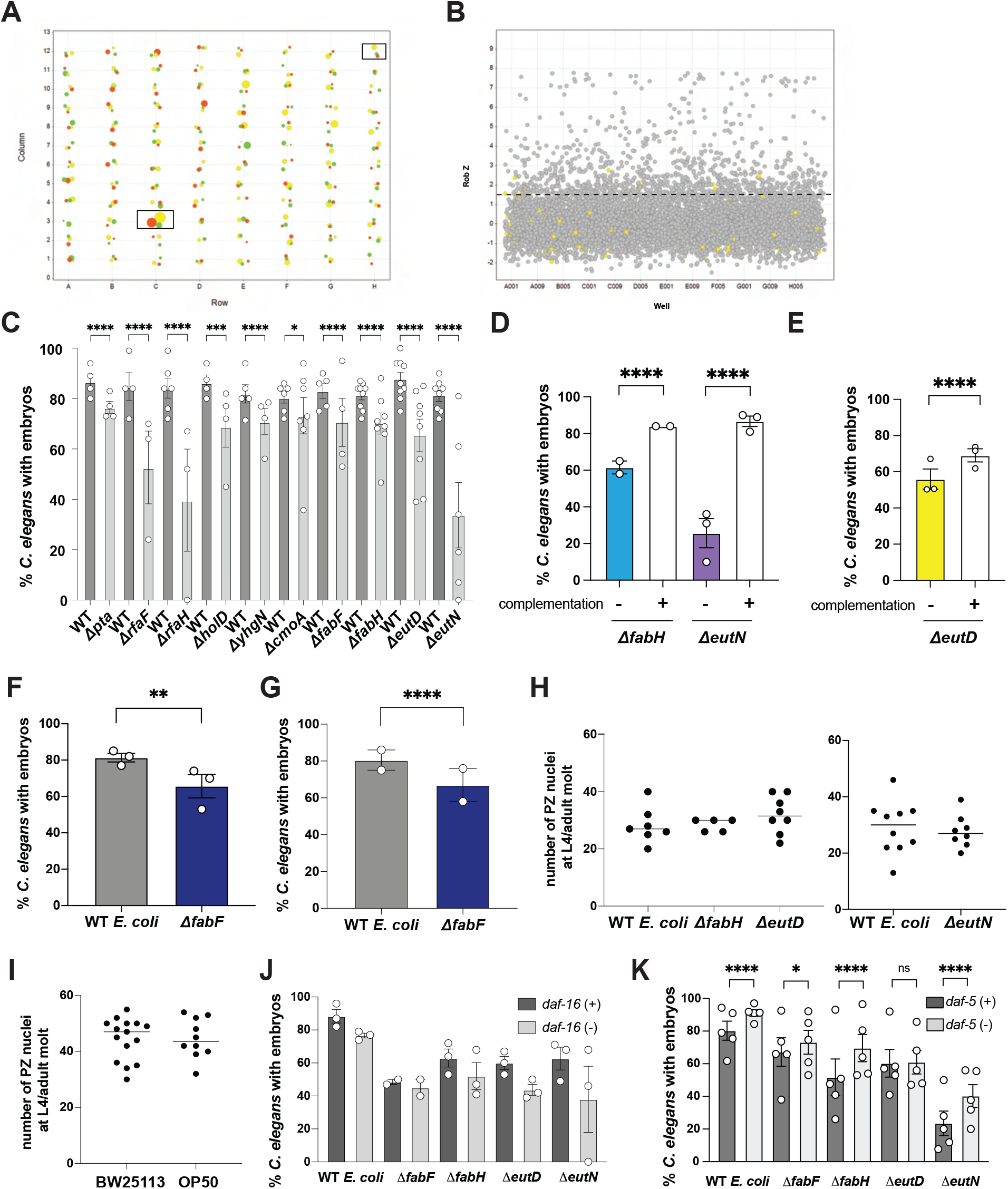
Primary screen selection criteria, validation of *E. coli* mutants, and effect of mutant *E. coli* on *C. elegans*, related to Fig. 1. **(A)** Representative bubble plot from secondary screen replicates. One of the six 96-well plates used in the secondary screen is shown here rendered using Vortex software data visualization tool. The size of the bubble represents percent of worms without embryos that is calculated from three technical replicates, while the color indicates the three biological trials. Boxed wells H12 and C3 indicate the negative control and a positive hit in two out of three replicates based on statistical tests. **(B)** Scatter plot showing the Robust Z (Rob Z) score for all primary screen wells scored. Scatter plot was generated using Vortex dotmatics software. Yellow dots indicate wells containing the negative control bacteria (GC1547). Dashed line indicates cut-off for Rob Z criterion. **(C)** Percentage of *C. elegans* bearing embryos in replicates of ten *E. coli* mutants identified from the secondary screen. Total n number of worms scored are (left to right): 505, 291, 418, 103, 879, 80, 586, 295, 474, 229, 1100, 501, 913, 411, 1253, 583, 1643, 550, 1561, 232. **(D)** Δ*eutN* and Δ*fabH* complementation suppresses *C. elegans* fertility defect. Total n number of worms scored is (left to right): 220, 197, 744, 1078. **(E)** Δ*eutD* complementation suppresses *C. elegans* fertility defect. Total n number of worms scored is 2276 (*ΔeutD*) and 2283 (*ΔeutD* with complementation). **(F, G)** Δ*fabF* phage transductant mimics *C. elegans* fertility defect of Δ*fabF* original clone. Circles represent mean of four independent biological replicates and bar is SEM. (F) Original *fabF* isolate from library (n = 330, 214 total worms scored, WT and *ΔfabF*, respectively), and (G) *fabF* transductant (n = 231, 66 total worms scored). **(H)** The number of progenitor zone nuclei of *glp-1(e2141)* at the L4 to Adult molt is not significantly different in worms raised on Δ*eutN*, Δ*fabH* or Δ*eutD* versus control *E. coli*. Scatter plots showing number of progenitor zone germ cell counts; each dot represents one gonad arm scored. We noted that in ∼30% of *eutN* mutants (relative to 10% in control), the border of the PZ was difficult to detect; these were censored from PZ counts. The elevated penetrance of this phenotype in the *ΔeutN*-fed worms may represent a meiotic entry defect, accounting for the delay in germline development relative to somatic development. **(I)** No significant difference is observed between the number of progenitors in worms grown on BW25113 versus OP50 on standard NGM solid media. We note that the number of progenitors is markedly lower in the *glp-1(e2141)* mutant background after growth on liquid versus growth on solid media; the Keio starting strain was used in (H) since OP50 is sensitive to kanamycin. **(J, K)** Fertility delay is not suppressed by loss of *daf-16,* nor is it unambiguously suppressed by loss of *daf-5*. Percent of worms with embryos on Day 1 in (J) *daf-16(+); daf-2(e1370) glp-1(e2141) or daf-16(m26); daf-2(e1370) glp-1(e2141)* and (K) *daf-5(+); glp-1(e2141) or daf-5(e1386); glp-1(e2141).* Loss of *daf-5* elevated the percentage of gravid worms on wild-type (WT) *E. coli*. Therefore, biological significance of a greater percentage of gravid *daf-5(e1386)* compared to *daf-5(+)* raised on the various mutant *E. coli* is unclear. Circles represent mean of independent biological replicates and bar is SEM. Total n number of worms scored are (left to right): **(J)** 269, 203, 207, 245, 179, 255, 229, 214, 200, 233 and (K) 460, 399, 386, 470, 379, 445, 388, 428, 554, 640. For (J), there is no significant suppression of the fertility delay in any case. Statistics: ANOVA and Student’s t-test (H and I, respectively) and Fisher’s Exact test (all other panels). * p < 0.05, ** p < 0.01, *** p < 0.001, **** p < 0.0001.

**Fig. S2.**
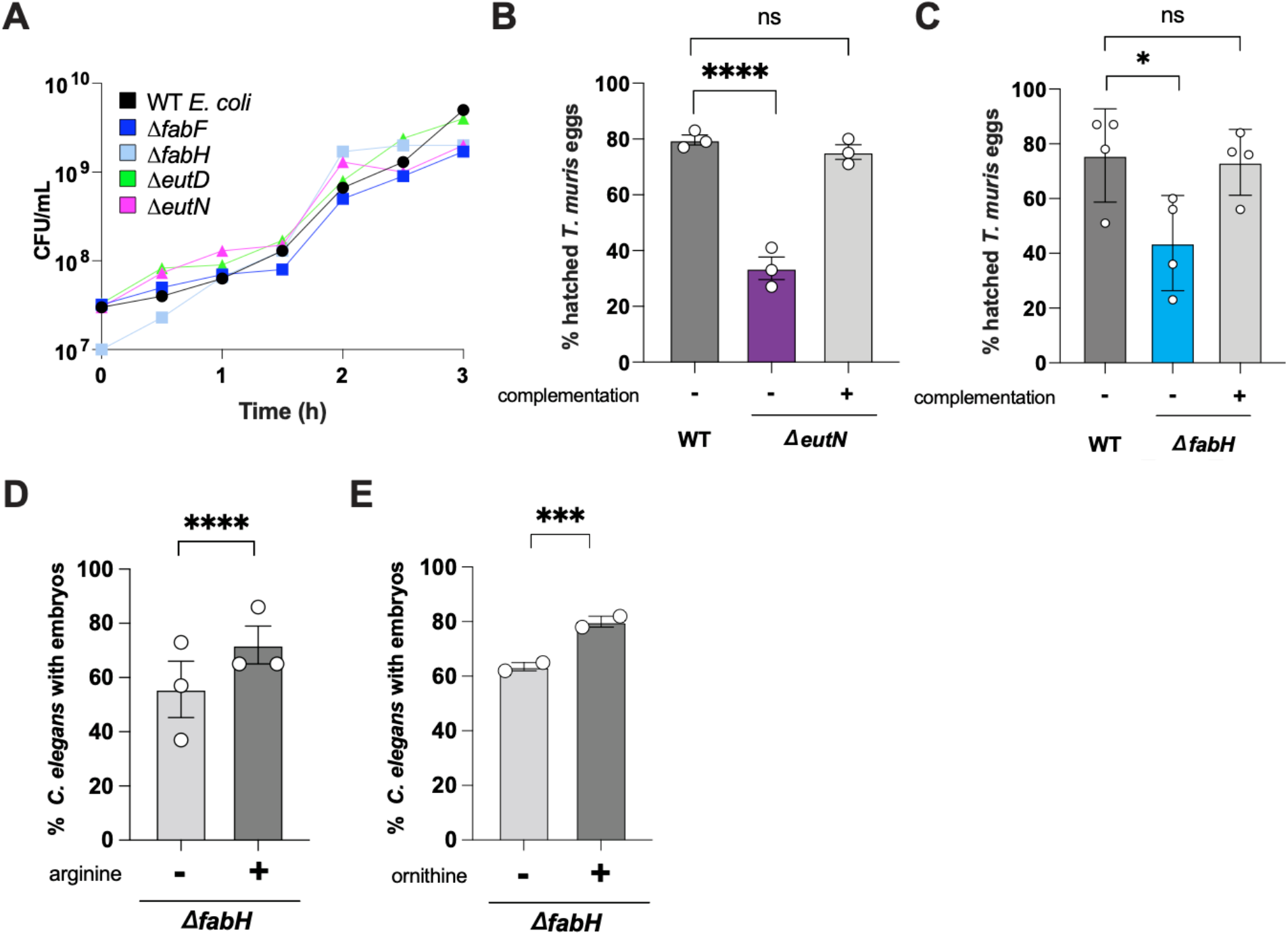
*E. coli* mutants used for *T. muris* egg hatching assay and effect of arginine and ornithine supplementation on *C. elegans* fertility, related to Fig. 2. **(A)** Bacterial density of monoclonal cultures of each of the strains indicated measured by dilution plating; CFU = colony forming units, h = hours. **(B-C)** Hatching elicited by WT *E. coli* (WT), mutants (B) Δ*eutN* and (C) Δ*fabH*, and strains where deletions were complemented with a plasmid-borne gene (n = 3-4). One-way analysis of variance (ANOVA) with Dunnett’s post-test compared with WT. **(D-E)** Percentage of *C. elegans* Day 1 adults with embryos raised on *ΔfabH E. coli* with arginine or ornithine supplementation. Fisher’s exact test (*ΔfabH* alone plus or minus supplementation with 500µM arginine or 100µM ornithine); Total n number of worms scored (left to right): (D) 237, 300 and (E) 216, 149. For all panels, measurements were taken from distinct samples. Data represent the average of at least two independent experiments. Graphs show means and SEM. * p < 0.05, *** p < 0.001, **** p < 0.0001. ns = not significant.

**Fig. S3.**
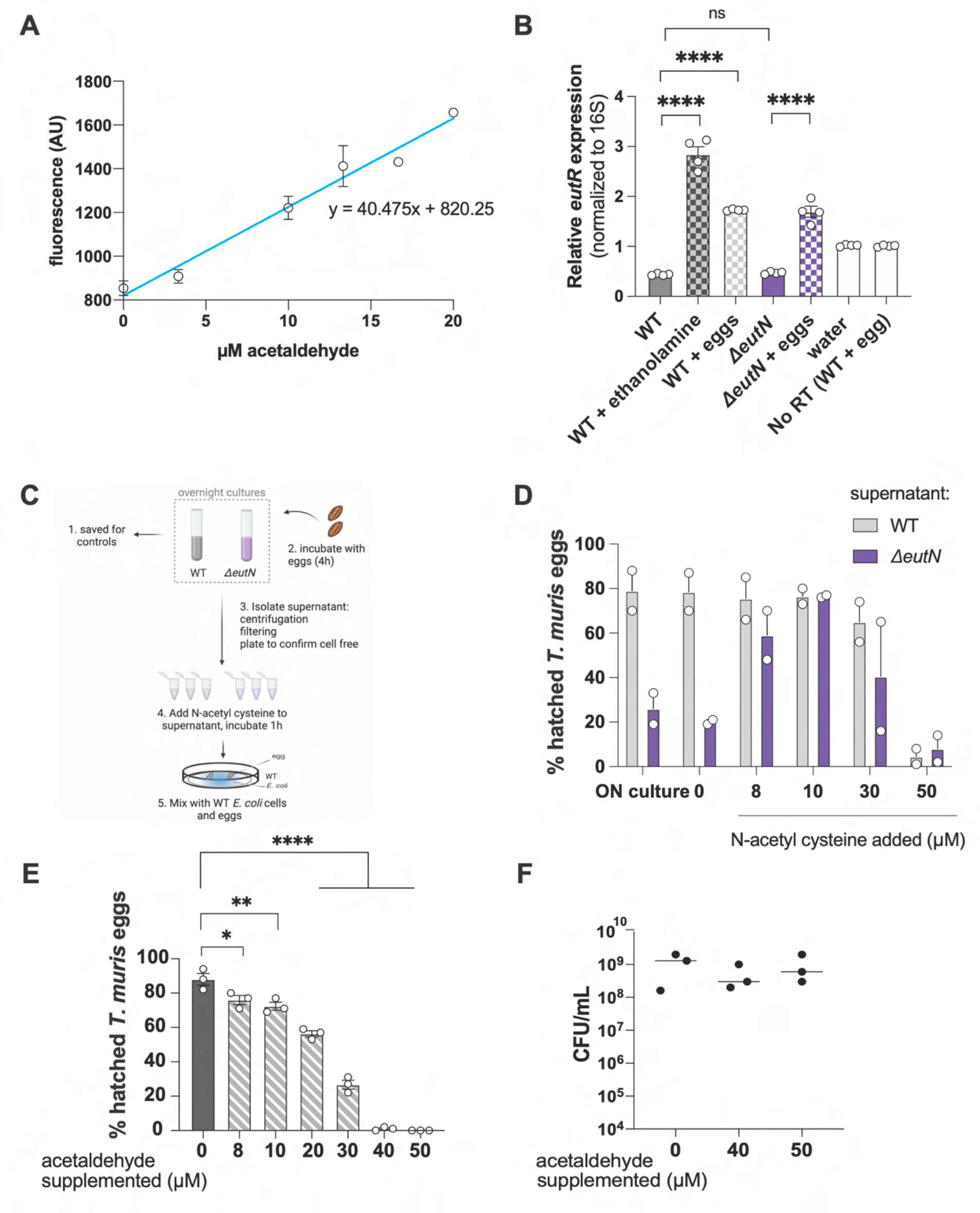
*E. coli*-derived and exogenous aldehydes reduce *T. muris* egg hatching, related to Fig. 3. **(A)** DarkZone standard curve for acetaldehyde. Fluorescence intensity of samples incubated with known concentrations of acetaldehyde and 20µM DarkZone dye (n = 2). AU, arbitrary units. **(B)** Relative *eutR* expression in *E. coli* cells. eggs = pre-incubation with *T. muris* eggs for 4 hours. No RT = no reverse transcriptase control. Data pooled from 2 independent experiments. Analyzed using ANOVA with Tukey’s multiple comparisons test. **(C)** Diagram of N-acetyl cysteine supplementation assay. Created using BioRender.com. **(D)** Quantification of hatching rates for mixtures containing WT *E. coli* cells and filtered supernatants supplemented with N-acetyl cysteine at the concentrations indicated (n = 2). **(E)** Quantification of hatching rates for *T. muris* eggs incubated with WT *E. coli* cultures supplemented with acetaldehyde at concentrations indicated (n = 3). Analyzed using ANOVA with Dunnett’s post-test compared to 0. **(F)** Bacterial density of conditions indicated from (E) measured by dilution plating (n = 3). For all panels, measurements were taken from distinct samples. Graphs show means and SEM. * p < 0.05, ** p < 0.01, **** p < 0.0001.

**Fig. S4.**
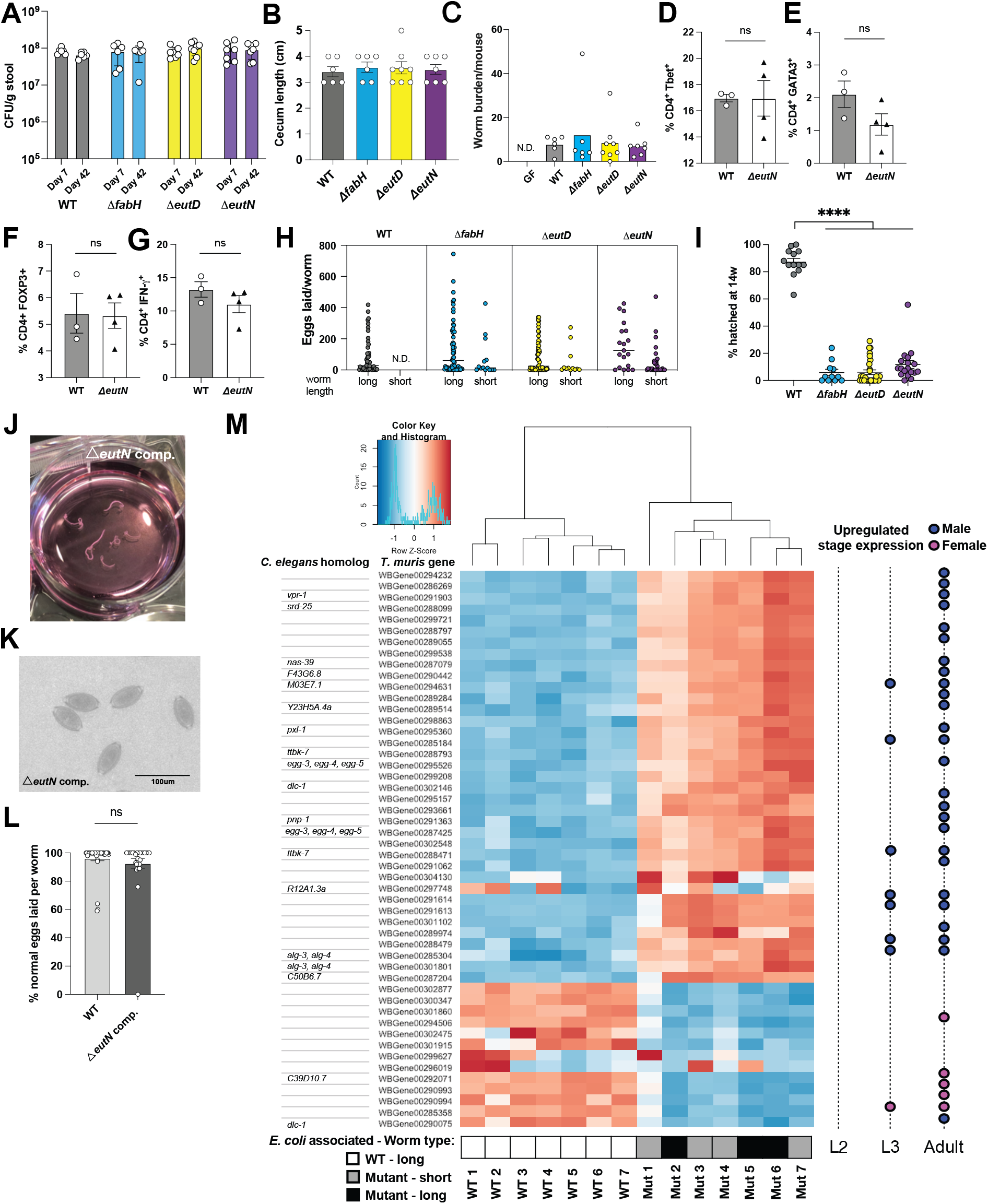
*T. muris* infection of germ-free and monocolonized mice, related to Fig 4. **(A)** *E. coli* burden 7 dpi and 42 dpi (day of sacrifice) in stool of mice gavaged with 1x 10^8^ CFU of strains indicated (n = 6-9 mice per group). **(B)** Cecum measurements taken on day of sacrifice for mice in (A). **(C)** Total worms harvested per mouse for germ-free (GF) and mice monocolonized with *E. coli* strains indicated; N.D. = not detected. **(D-G)** CD4^+^ T cell percentage from mesenteric lymph nodes of WT and *ΔeutN* monocolonized mice. (D) Th1 (Tbet^+^), (E) Th2 (GATA3^+^), (F) Treg (FOXP3^+^). (G) Cells were also stimulated with PMA and ionomycin and stained for intracellular IFN-γ expression, a cytokine known to be positively correlated with host susceptibility to *T. muris*. **(H)** Quantification of eggs laid per worm disaggregated by worm length. Short worms measured < 1 cm. N.D. = not detected. Harvested worms were incubated in individual wells of a 48-well plate with RPMI medium overnight and eggs laid were quantified using light microscopy. **(I)** Percentage of eggs subjected to 14 weeks of embryonation that hatched after incubation with WT *E. coli*. **(J)** Representative image of worms harvested from mice monocolonized with *ΔeutN* strain where *eutN* has been complemented on a plasmid = *ΔeutN* comp. (WT, n = 9; *ΔeutN* comp., n = 8). **(K)** Representative image of eggs laid by worms harvested from mice monocolonized with *ΔeutN* comp. **(L)** Proportion of total eggs laid per worm with normal morphology. Circles represent individual worms (WT, n = 36; *ΔeutN* comp., n = 29). **(M)** Heatmap of the 50 most differentially detected genes between WT- and mutant-associated worms generated using DESeq2. Colors indicate Row Z-score and clusters of genes with similar trends in expression between samples. Red indicates upregulated genes and blue indicates downregulated genes. WT = wildtype *E. coli*. L = *T. muris* larval stage. For all panels, measurements were taken from distinct samples. Data pooled from two independent experiments. Graph shows means and SEM. **** p < 0.0001.

**Table S1.** Top candidates and bacterial gene function (EcoCyc). *Fab* and *eut* genes were chosen for further characterization.

| **GENE** | **PROTEIN FUNCTION** | **FUNCTIONAL GROUP** |
| --- | --- | --- |
| *fabF* | β-ketoacyl-[acyl carrier protein] synthase II | Fatty acid biosynthesis |
| *fabH* | β-ketoacyl-[acyl carrier protein] synthase III | Fatty acid biosynthesis |
| *eutD* | phosphate acetyltransferase | Ethanolamine utilization |
| *eutN* | putative ethanolamine catabolic microcompartment shell protein | Ethanolamine utilization |
| *pta* | phosphate acetyltransferase | Ethanolamine utilization |
| *rfaF* | ADP-heptose—LPS heptosyltransferase 2 | Lipopolysaccharide biosynthesis |
| *rfaH* | transcription antiterminator RfaH | Lipopolysaccharide biosynthesis |
| *cmoA* | carboxy-S-adenosyl-L-methionine synthase | Uridine modification in RNA |
| *yhgN* | putative inner membrane protein | Antibiotic resistance |
| *holD* | DNA polymerase III subunit $\psi$ | DNA replication |

**Table S2.**
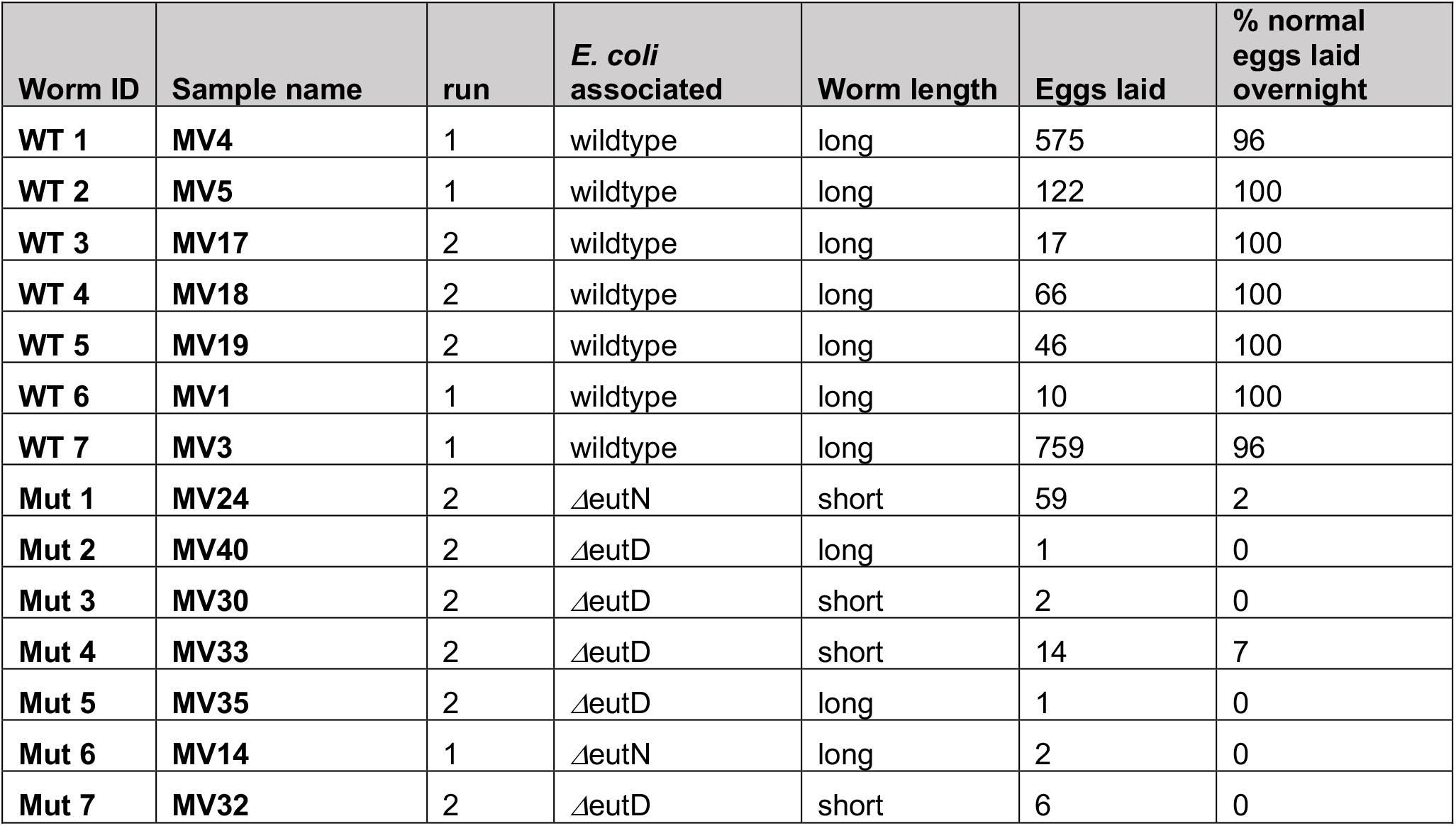
Single-worm Seq sample metadata, related to Figures 4 and S4. Data for *T. muris* worms subjected to RNA extraction and sequencing analysis.

**Table S3.**
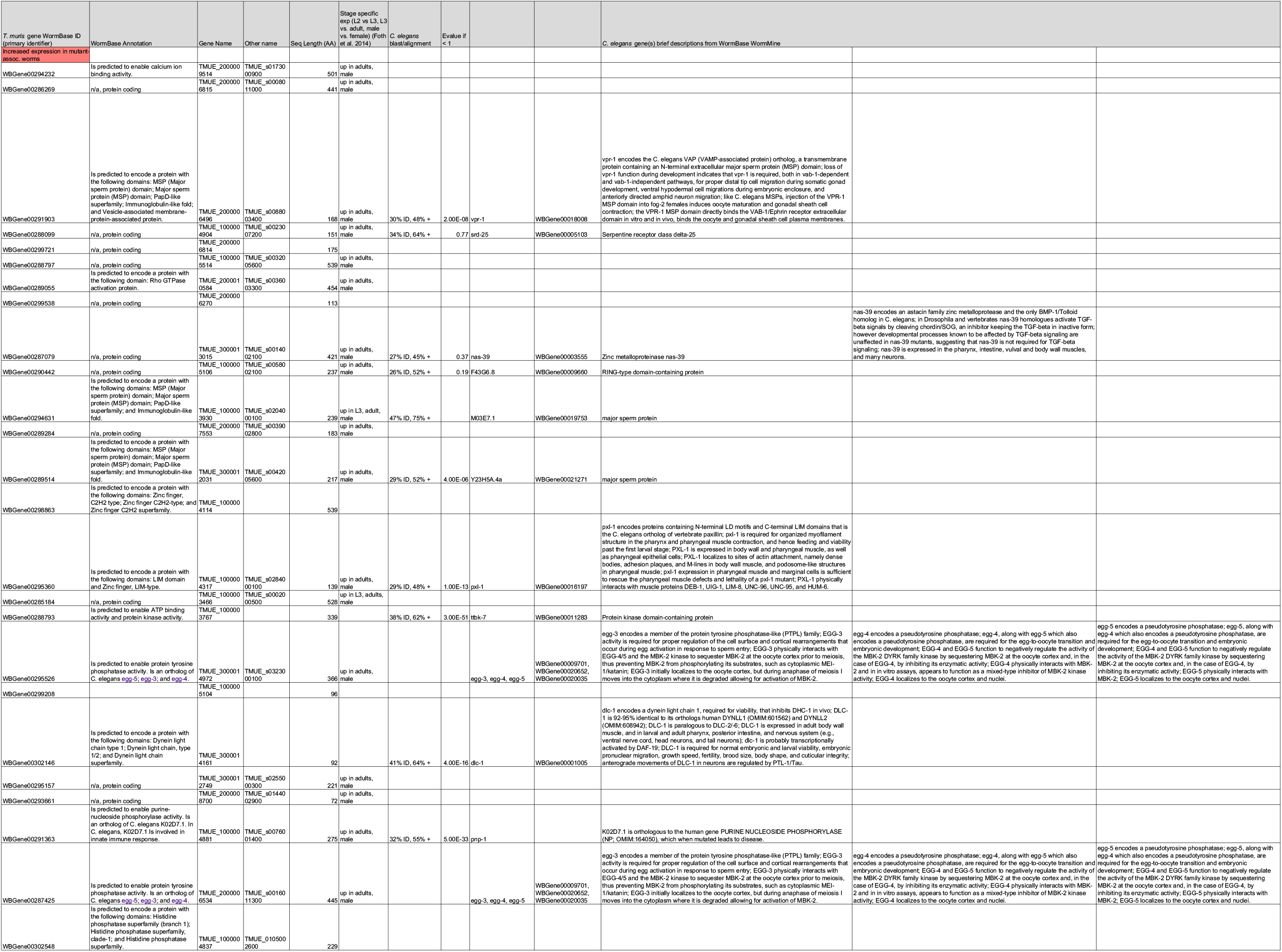
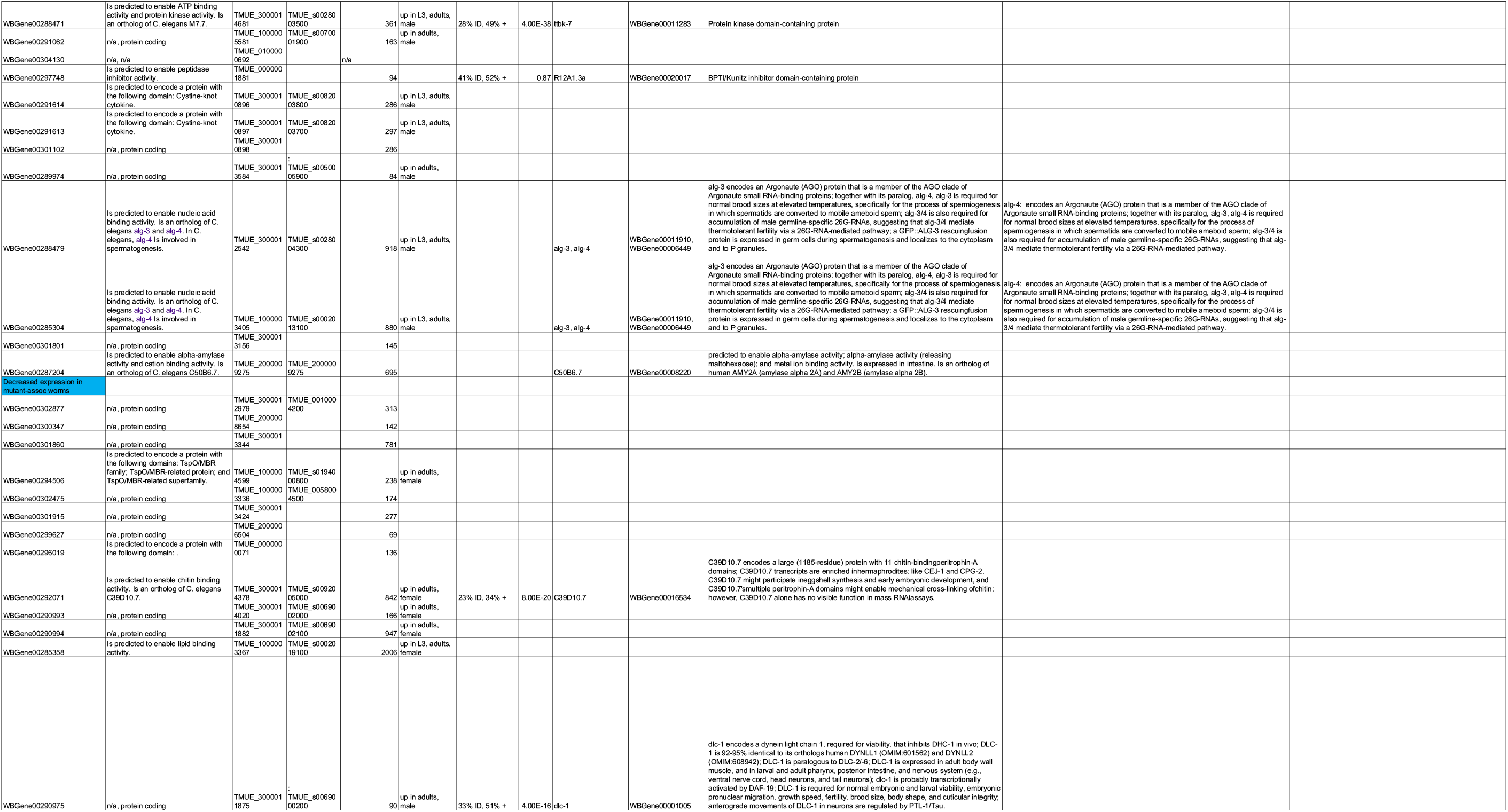
List of top 50 differentially detected genes, related to Figure S4E. Information was collated from WormBase (version WS283) and Foth et al. (2014) for stage specific expression data*. C. elegans* homologs were found using WormBase BLAST/BLAT.

**Table S4.**
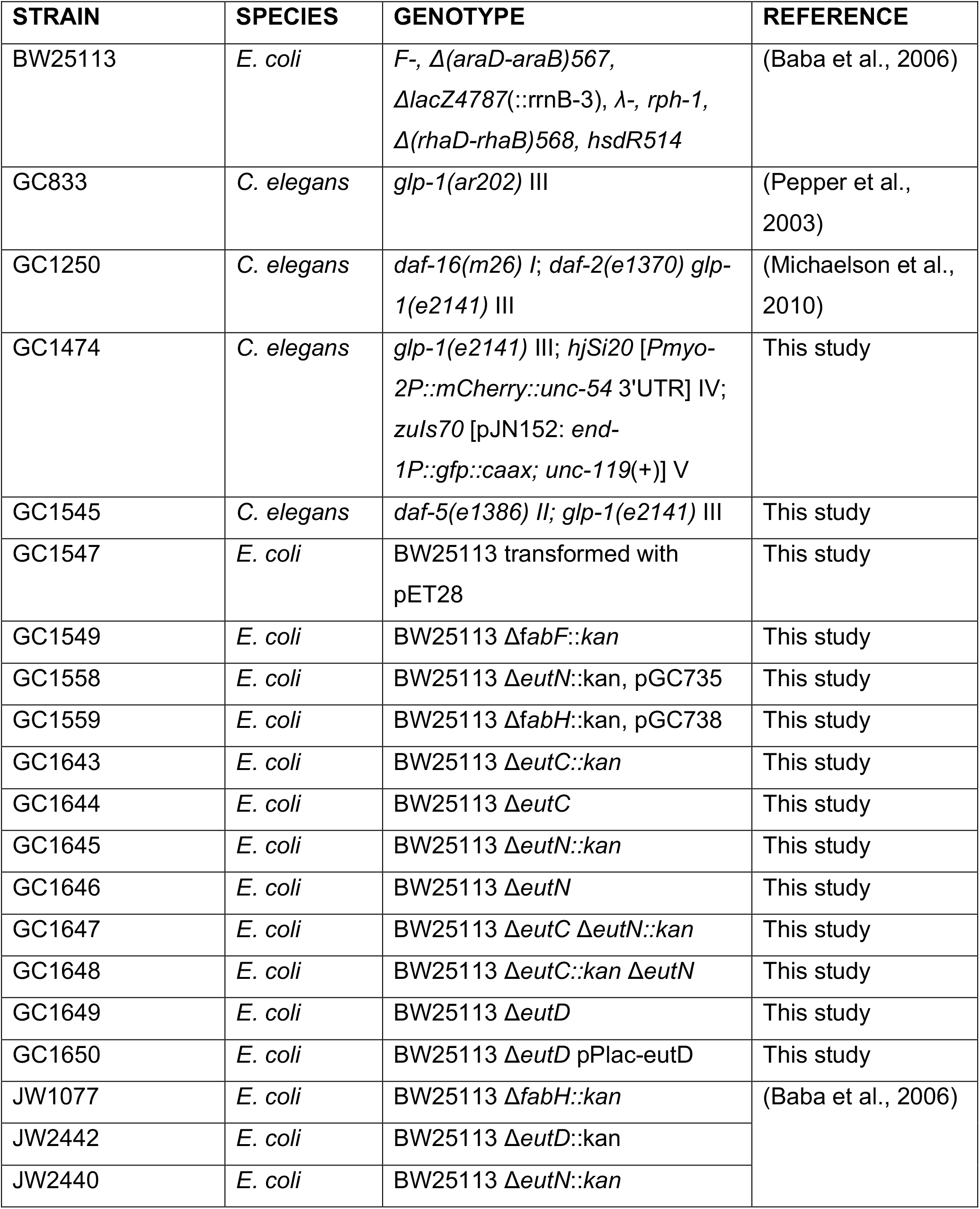
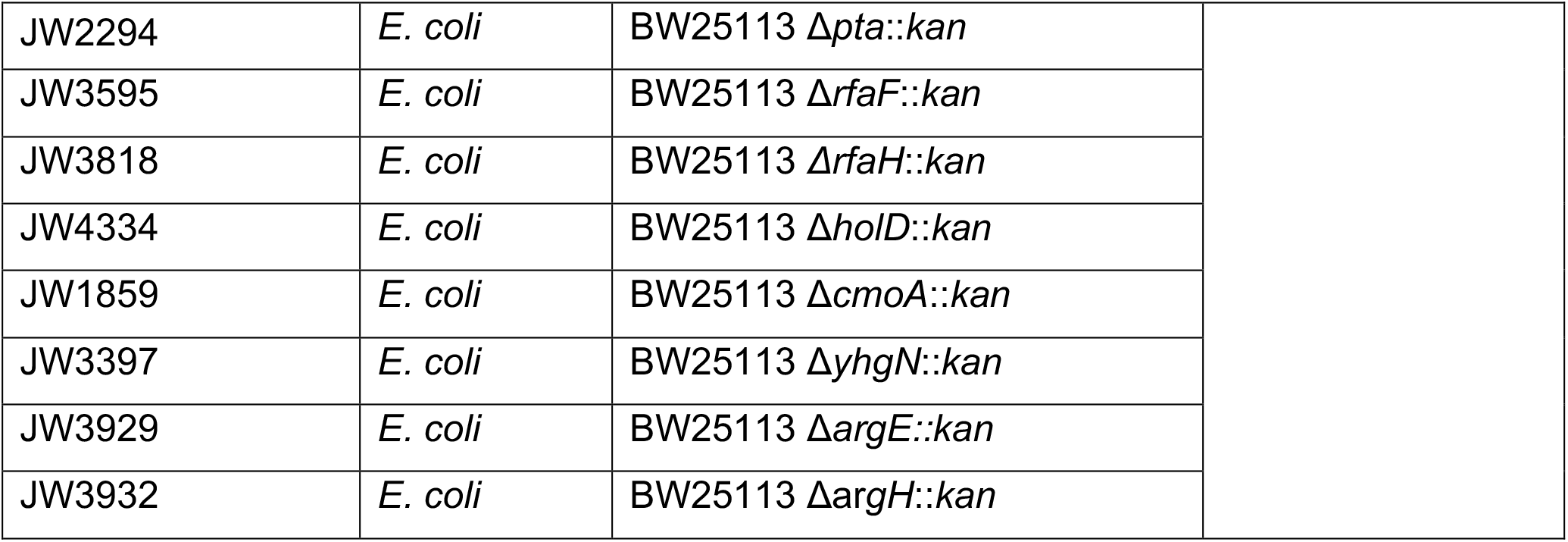
Strains used in this study.

| STRAIN | SPECIES | GENOTYPE | REFERENCE |
| --- | --- | --- | --- |
| BW25113 | *E. coli* | *F*-, $\Delta(araD-araB)567$, $\Delta lacZ4787(::rrnB-3)$, λ-, *rph-1*, $\Delta(rhaD-rhaB)568$, *hsdR514* | (Baba et al., 2006) |
| GC833 | *C. elegans* | *glp-1(ar202)* III | (Pepper et al., 2003) |
| GC1250 | *C. elegans* | *daf-16(m26)* I; *daf-2(e1370)* *glp-1(e2141)* III | (Michaelson et al., 2010) |
| GC1474 | *C. elegans* | *glp-1(e2141)* III; *hjSi20* [*Pmyo-2P::mCherry::unc-54* 3'UTR] IV; *zuls70* [pJN152: *end-1P::gfp::caax*; *unc-119(+)*] V | This study |
| GC1545 | *C. elegans* | *daf-5(e1386)* II; *glp-1(e2141)* III | This study |
| GC1547 | *E. coli* | BW25113 transformed with pET28 | This study |
| GC1549 | *E. coli* | BW25113 $\Delta fabF::kan$ | This study |
| GC1558 | *E. coli* | BW25113 $\Delta eutN::kan$, pGC735 | This study |
| GC1559 | *E. coli* | BW25113 $\Delta fabH::kan$, pGC738 | This study |
| GC1643 | *E. coli* | BW25113 $\Delta eutC::kan$ | This study |
| GC1644 | *E. coli* | BW25113 $\Delta eutC$ | This study |
| GC1645 | *E. coli* | BW25113 $\Delta eutN::kan$ | This study |
| GC1646 | *E. coli* | BW25113 $\Delta eutN$ | This study |
| GC1647 | *E. coli* | BW25113 $\Delta eutC \Delta eutN::kan$ | This study |
| GC1648 | *E. coli* | BW25113 $\Delta eutC::kan \Delta eutN$ | This study |
| GC1649 | *E. coli* | BW25113 $\Delta eutD$ | This study |
| GC1650 | *E. coli* | BW25113 $\Delta eutD$ pPlac-eutD | This study |
| JW1077 | *E. coli* | BW25113 $\Delta fabH::kan$ | (Baba et al., 2006) |
| JW2442 | *E. coli* | BW25113 $\Delta eutD::kan$ | |
| JW2440 | *E. coli* | BW25113 $\Delta eutN::kan$ | |
| JW2294 | *E. coli* | BW25113 Δ*pta::kan* | |
| JW3595 | *E. coli* | BW25113 Δ*rfaF::kan* | |
| JW3818 | *E. coli* | BW25113 Δ*rfaH::kan* | |
| JW4334 | *E. coli* | BW25113 Δ*holD::kan* | |
| JW1859 | *E. coli* | BW25113 Δ*cmoA::kan* | |
| JW3397 | *E. coli* | BW25113 Δ*yhgN::kan* | |
| JW3929 | *E. coli* | BW25113 Δ*argE::kan* | |
| JW3932 | *E. coli* | BW25113 Δ*argH::kan* | |

**Table S5.**
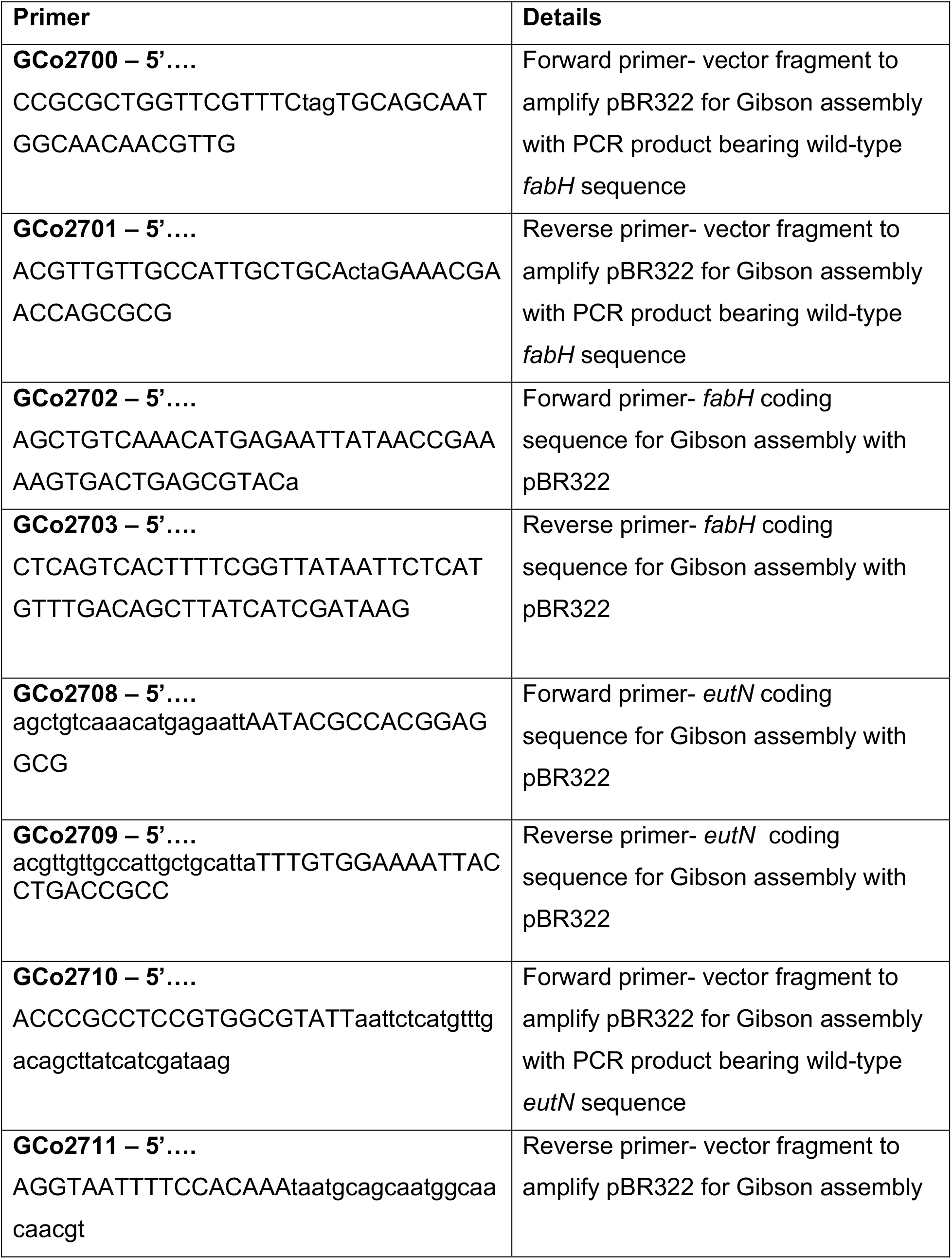
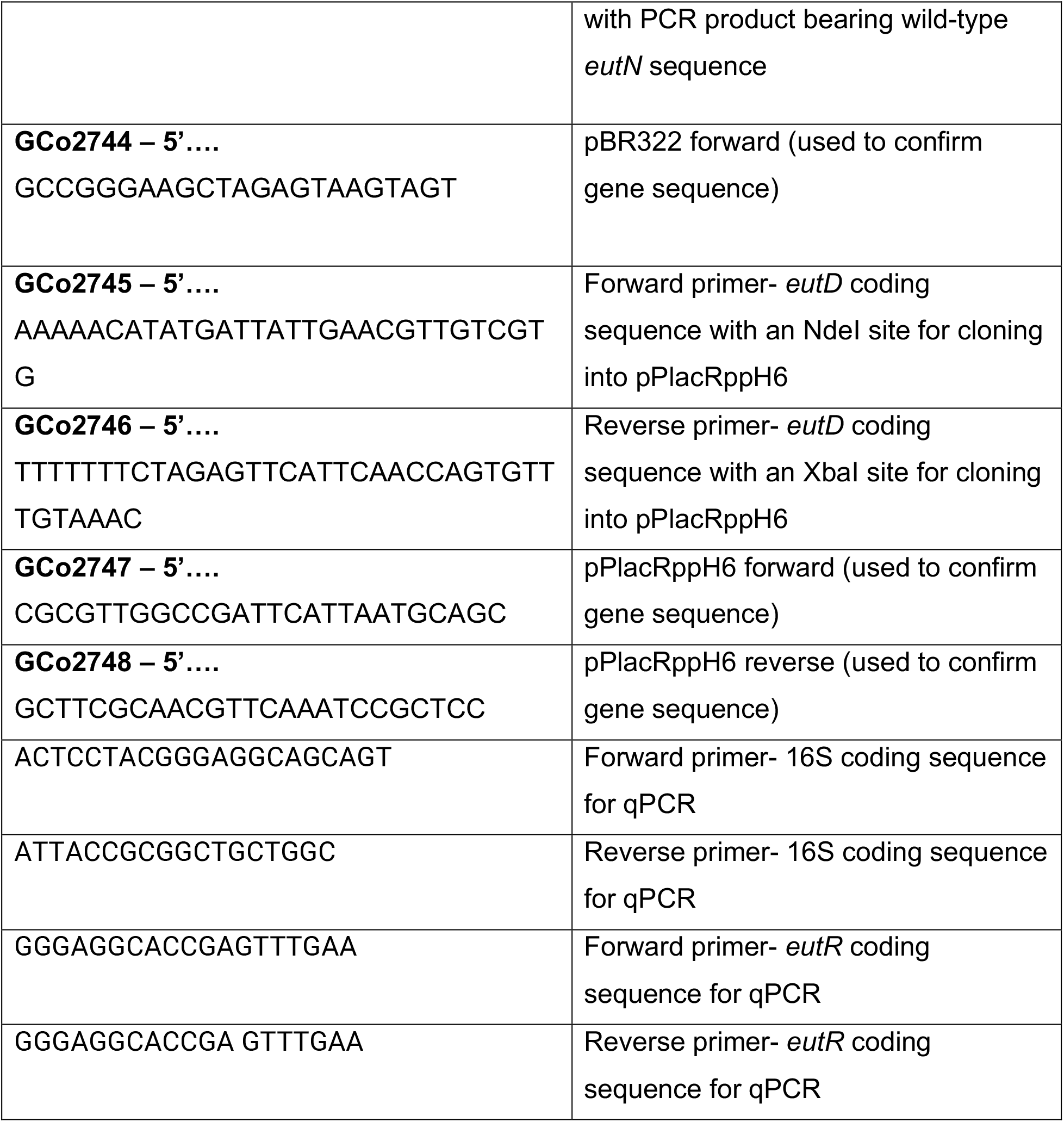
Primers used in this study.

**Table S6.** Plasmid names and descriptions.

| Plasmid name | Description |
| --- | --- |
| pGC735 | *eutN* cloned into pBR322 by Gibson assembly |
| pGC737 | pET28 |
| pGC738 | *fabH* cloned into pBR322 by Gibson assembly |
| pPlac-eutD | *eutD* under the control of a *lacUV5* promoter |
| pBR322 | pBR322 |

**Table S7.**
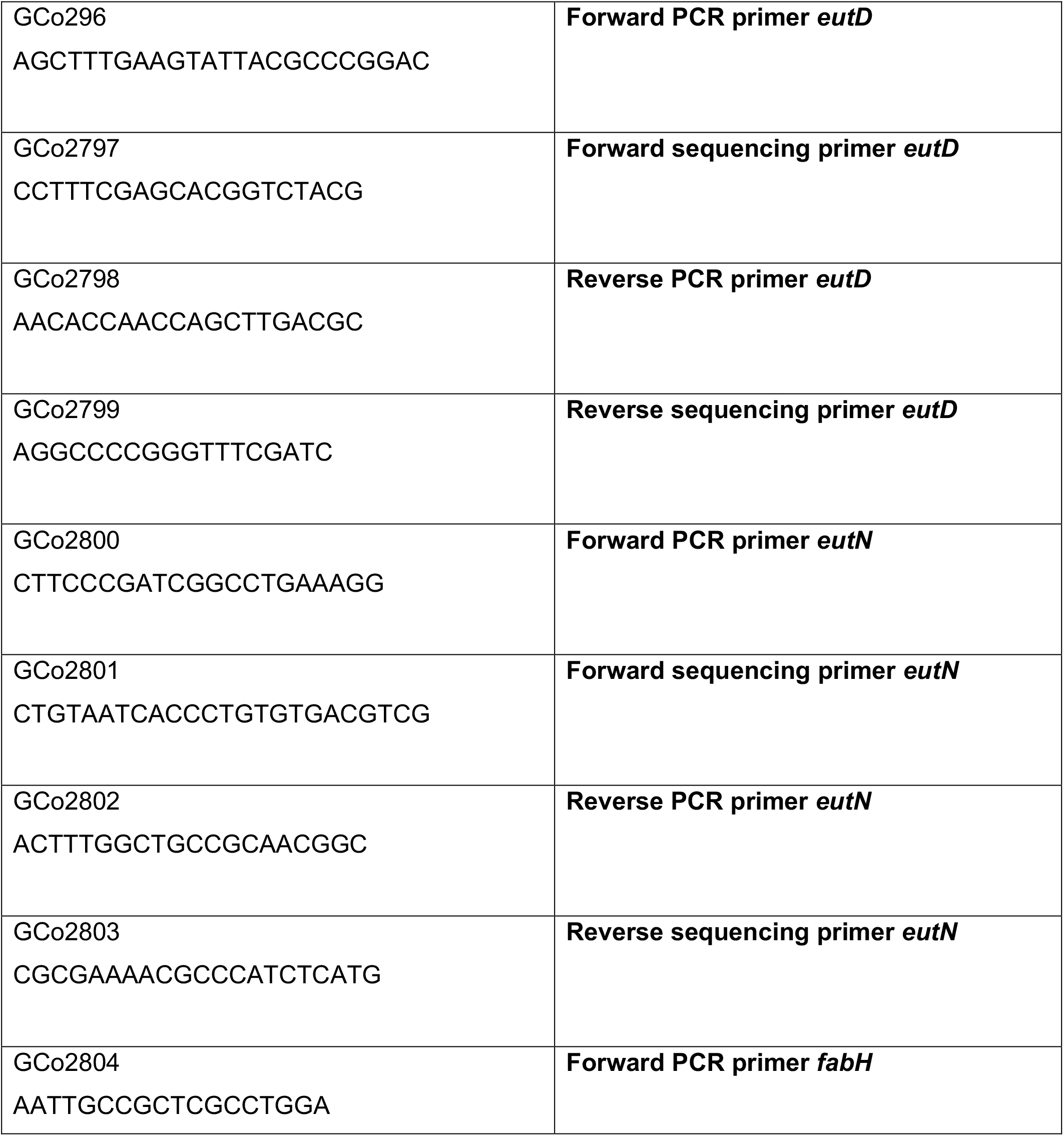
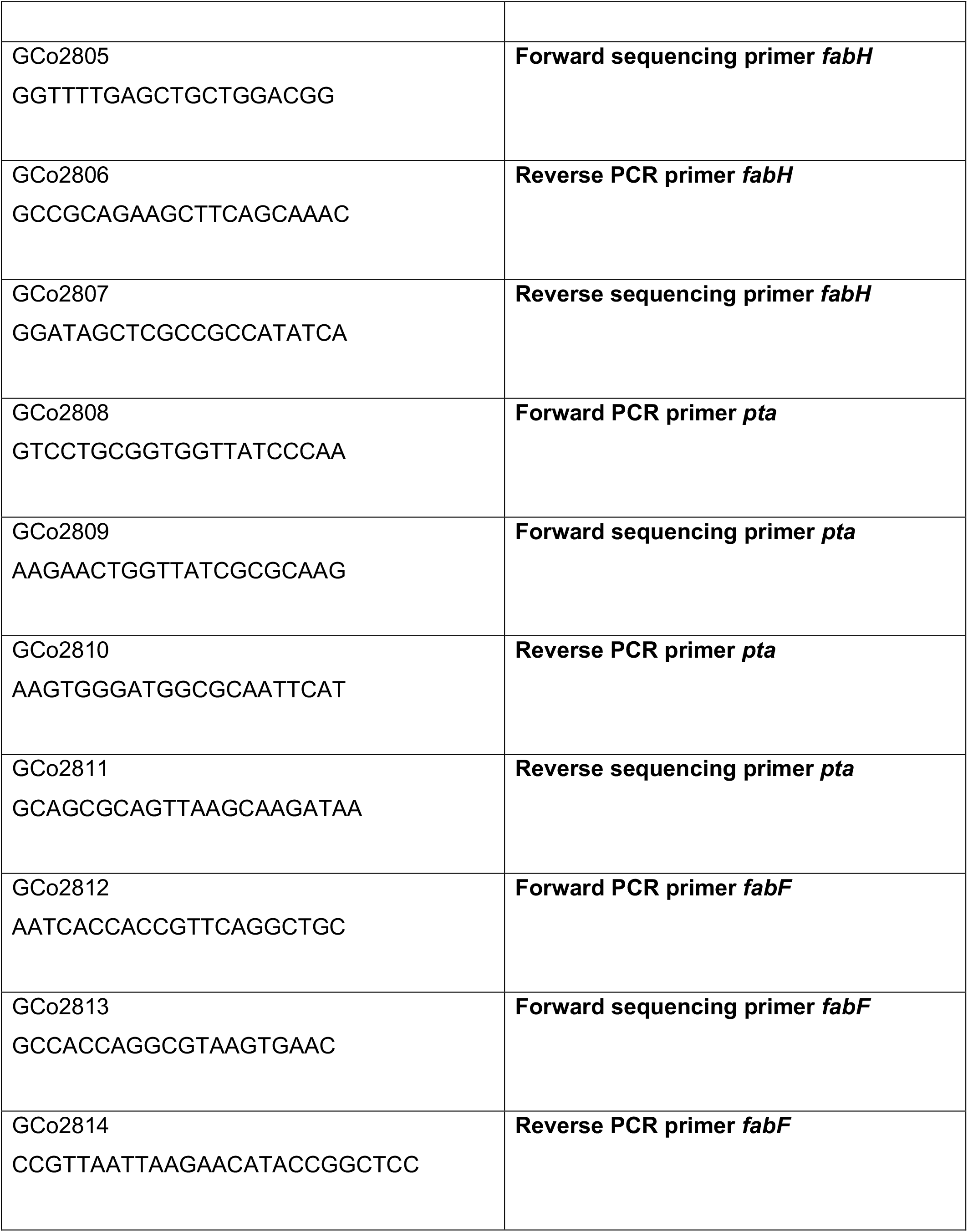
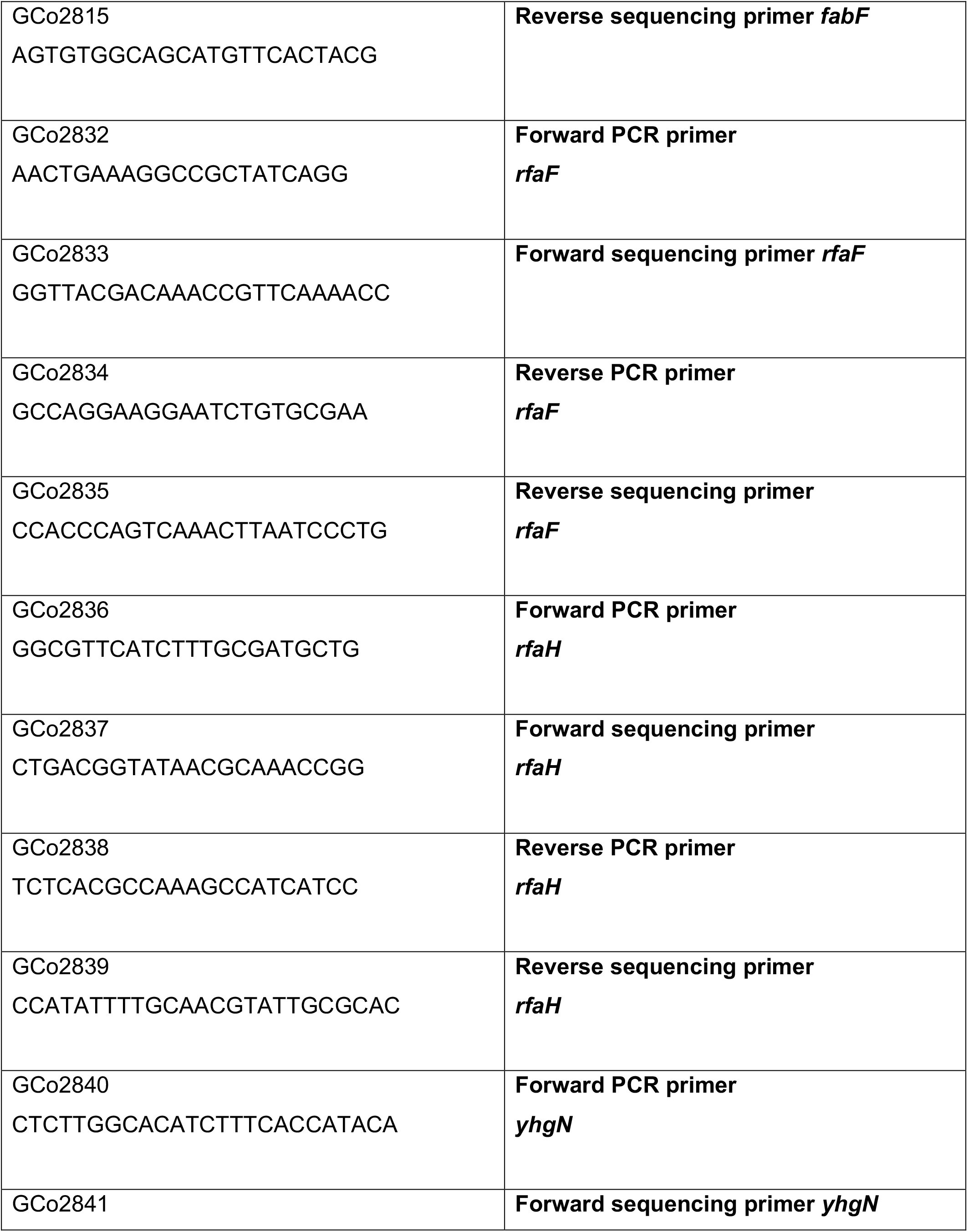
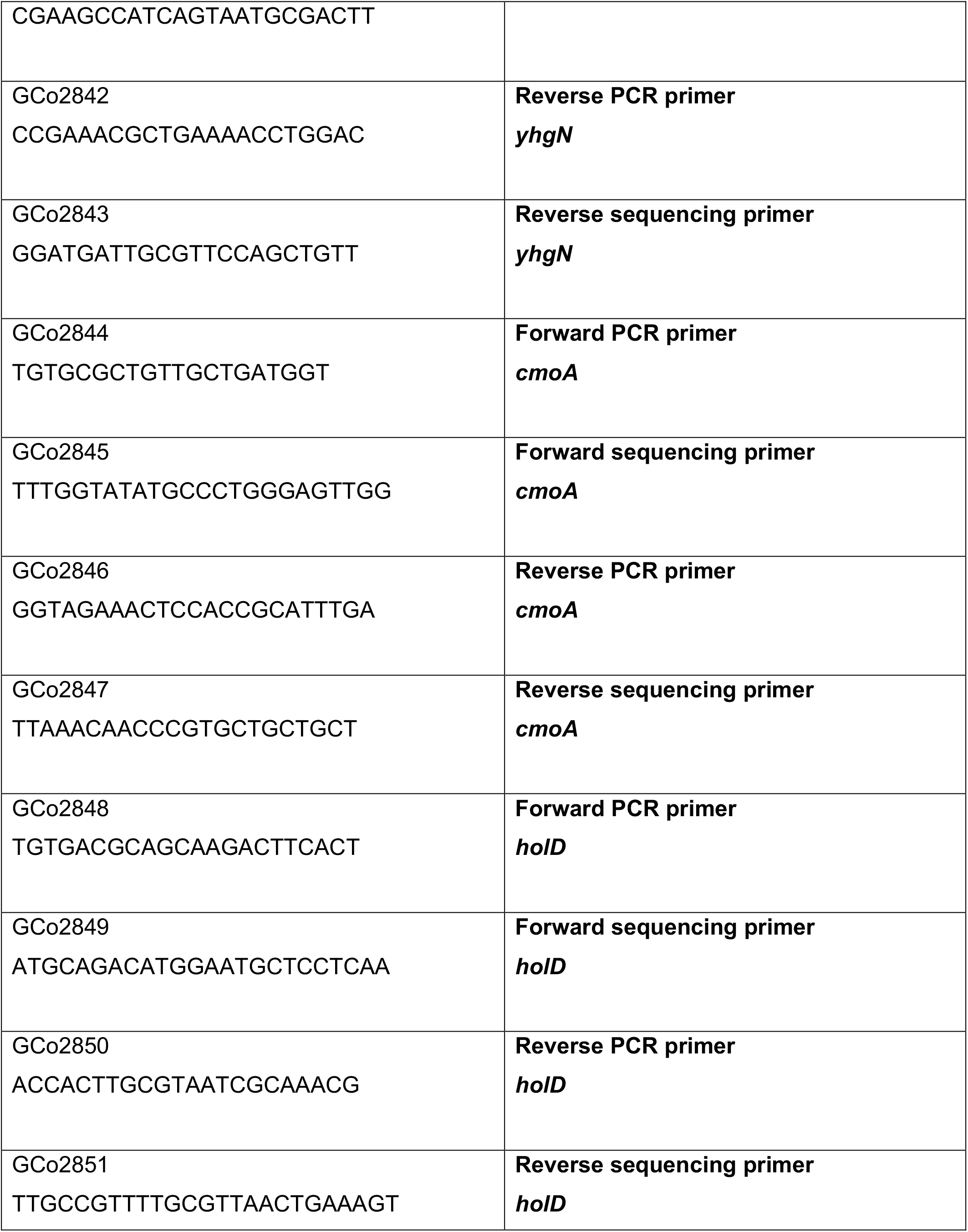
Primer sequences to verify kan cassette insertion within target genes.

